# Cheminformatic identification of small molecules targeting acute myeloid leukemia

**DOI:** 10.1101/2025.05.20.655224

**Authors:** Megan R. Daneman, Bernadetta Meika, Lois Armendariz, Elissa Tjahjono, Alexey V. Revtovich, Leonid A. Stolbov, Natthakan Thongon, Natalia Baran, Scott R. Gilbertson, Vladimir V. Poroikov, Natalia V. Kirienko

## Abstract

Acute myeloid leukemia (AML) is an aggressive hematological malignancy with poor prognosis and high relapse rates when treated with cytotoxic chemotherapeutics. Previously, we identified a family of small molecules that modulate mitochondrial function, referred to as PS127-family compounds. These drugs were selectively toxic to AML and were characterized by two predicted functions: apoptotic agonism and thioredoxin/glutathione reductase inhibition. Here, we uncovered a third critical predicted function, autophagic induction. Using a cheminformatic screen of ∼4.2 million compounds for molecules with high predicted probability for all three functions, we found and validated hits that selectively killed AML cells, activated apoptosis, were dependent upon autophagic activation, and compromised glutathione metabolism by interfering with glutathione reductase, all of which are consistent with predictions. Compound treatment increased pools of cytosolic and mitochondrial ROS, decreased oxygen consumption, and reduced ATP synthesis. Structurally-unrelated compounds caused the same phenotypes, validating our approach of screening for predicted function. Finally, we also observed strong synergy between these compounds and midostaurin and venetoclax, underscoring their therapeutic potential. Key phenotypes, including the compounds’ impact on glutathione metabolism and synergy with doxorubicin and midostaurin, were confirmed in AML-patient-derived primary cells, validating the potential of these compounds for the development of future AML treatments.

## Introduction

Acute myeloid leukemia (AML) is an aggressive, heterogeneous, hematopoietic malignancy of myeloid origin that can be rapidly fatal if left untreated [1]. Treatment of AML is complicated by the presence of two very distinct populations of cancer cells: leukemic stem cells and blast cells. Bulk leukemic myeloblasts are numerous, incompletely differentiated, and often immunologically incompetent. In AML patients, blast cells are produced from precursor cells at an increased rate relative to normal hematopoiesis and, in some cases, cells become so numerous that they can cause leukostasis and organ failure [2]. This population is targeted by the most common first-line AML treatment regimen, called induction and consolidation, which consists of two foundational drugs: the pyrimidine analog cytosine arabinoside (cytarabine; ara-C) and an anthracycline drug, given in combination. While this therapy is usually effective at targeting and eliminating rapidly dividing bulk cell populations, reducing the number of circulating myeloblasts [3].

This approach fails at eliminating leukemic stem cells (LSCs), which are more slowly cycling and protected in the bone marrow, where drug exposure is reduced [4]. These cells are thought to be a primary driver of relapse, as they continually give rise to new cancerous precursor cells that will begin to divide more rapidly and replenish the blast population [5]. Worse, induction and consolidation may select for treatment-resistant clones, further complicating treatment [4]. Effective AML therapy requires strategies capable of targeting both.

Notably, recent studies have begun to reveal that both cell populations share metabolic vulnerabilities that can be targeted for treatment [6,7]. For example, AML cells depend more heavily on oxidative phosphorylation (OXPHOS) and glutaminolysis to meet their energetic needs compared to normal hematopoietic stem and progenitor cells [4,8–11]. Previously, we discovered that mitochondria in AML cells also have lower coupling efficiency, likely due to increased proton leakage [12]. Increased OXPHOS activity drives correspondingly higher production of reactive oxygen species (ROS), which damage cellular macromolecules including lipids, proteins, and nucleic acids in the cell [13,14]. Mitochondria, the primary cellular source of ROS, are themselves highly-susceptible to ROS-induced damage[6], which impairs their function and efficiency, leading to further ROS generation and establishing a self-perpetuating cycle of oxidative stress.

Mitochondria have limited intrinsic repair pathways; one of the most important of these is mitophagy – the selective autophagy recycling of damaged mitochondria. This process is heavily regulated for homeostatic maintenance, as both insufficient and excessive mitophagy can trigger programmed cell death [15]. For this reason, mitophagy has been increasingly investigated as a potential method for inducing cancer cell-specific death over the past decade with mixed results, as both the induction and inhibition of mitophagy can induce leukemia cell death [4,8,16–20].

We previously showed that AML cells are very sensitive to combinations of different mitochondria-targeting drugs with established AML agents, including IACS-10759 and vinorelbine [21]. We also showed that several novel activators of PINK1-mediated mitophagy selectively killed AML cells [12]. These PINK1-stabilizing (PS) compounds interfered with mitochondrial bioenergetics, characterized by decreased routine and ATP-linked respiration, and increased mitochondrial membrane depolarization—ultimately leading to cell death in blasts and LSCs. The most effective compounds identified, PS127E and PS127B, were chemically similar derivatives anchored by a parent lead compound, PS127, identified in a high-content phenotypic screen in *Caenorhabditis elegans* [22]. Cheminformatic analysis revealed that PS127, PS127B, and PS127E were likely inhibitors of key enzymes governing redox homeostasis, specifically thioredoxin reductase and glutathione reductase [12].

In this study, we used a ligand-based, cheminformatic approach to screen over 4.2 million compounds to identify small molecules with the same predicted functions as the PS127 family (apoptotic agonism and thioredoxin or glutathione reductase (TrxR and/or GR) inhibition to which we also added autophagic induction). This screen identified 93 hits, of which 81 were structurally similar to the PS127 family. These compounds exhibited synergy with approved AML therapeutics, particularly midostaurin (MID), indicating potential as combinatorial treatments for AML. To uncover their mechanism of action, we verified three predicted activities for selected PS127 analogs. These compounds triggered apoptosis, required functional autophagy for AML cell cytotoxicity, and disrupted glutathione metabolism. Importantly, we demonstrated direct interaction between the drugs and glutathione reductase, validating it as the molecular target. Notably, a structurally-unrelated compound was also predicted to have the same three activities and demonstrated similar experimental outcomes, corroborating our approach. These compounds compromised mitochondrial metabolism in AML cells, reducing oxygen consumption rates (OCR) and ATP levels. These findings demonstrate the utility of *in silico,* function-based cheminformatics screening anchored to known bioactive compounds for the identification of novel drug leads and uncovered the mechanism of action for PS127-family compounds.

## METHODS

### Cell lines, progenitor cells, and patient samples

AML cell lines were purchased from ATCC (Manassas, VA, USA). Cryopreserved AML patient samples collected between 2018 and 2019 were supplied by MD Anderson Cancer Center Leukemia Sample Bank **(Supplementary Table 1)**. Cell culturing was performed as previously described [21], and peripheral blood mononuclear cells (PBMC) were isolated as described therein. Human bone marrow-derived CD34^+^ cells were purchased from AllCells, a Discovery Life Sciences Company (Alameda, CA, USA) (**Supplementary Table 2**). Further details on cell culturing are described in **Supplementary Methods 1**.

### In silico screen

Cheminformatics screening was performed using Prediction of Activity Spectra for Substances (PASS) version 2022 [23]. Screened molecules were clustered based on structural similarity using Multidimensional Scaling (MDS) in ChemMine Tools (cut off ≥ 0.4) [24]. Structural similarity was analyzed further using the Tanimoto coefficient based on PubChem fingerprinting and visualized in Cytoscape 3.10.3. Further details are outlined in **Supplementary Methods 2**.

### Cytotoxicity assays

All compounds identified *in silico* were purchased from Molport. Cells were treated with compounds or solvent controls in experimental media (RPMI-1640 with 1% FBS, 1% P/S) and stained with Hoechst 33342 and propidium iodide (PI) to determine total cell count and dead cells, respectively, as previously described [25]. Human bone marrow-derived CD34^+^ cells were treated in experimental media (StemSpan SFEM II with 1% penicillin/streptomycin (P/S) and glutamine). Further details on single drug cytotoxicity assay and drug synergy are outlined in **Supplementary Methods 3**.

### Thioredoxin reductase and glutathione reductase assays

Thioredoxin reductase (TrxR) activity in MOLM-13 cells was assessed using the Thioredoxin Reductase Colorimetric Assay Kit (Cayman Chemicals) and compound-mediated disruption of glutathione redox balance (glutathione reductase (GR) interference, understood as the ratio of reduced glutathione (GSH) to oxidized glutathione (GSSG)) were evaluated using the EnzyChrom™ Glutathione GSH/GSSG Assay Kit according to the respective manufacturer’s instructions. GR activity in cell lysates was measured using DetectX® Glutathione Reductase Fluorescence Activity Kit (Arbor Assay, Michigan) according to the manufacturer’s instructions. The physical interaction between compounds of interest and GR was evaluated via differential scanning fluorometry (DSF), where binding was indicated by a shift in the protein melting temperature. Further details are outlined in **Supplementary Method 4 and 5**.

### Measurements of autophagy

Autophagy was assessed by co-treating MOLM-13 cells with known early and late autophagic inhibitors using the drug combination protocol described in **Supplementary Methods 3**. Early- stage autophagy inhibitors tested included wortmannin (TargetMol Chemical) and 3-methyladenine (3-MA) (TCI America™). Late-stage autophagic inhibitors tested were hydroxychloroquine (HCQ) and chloroquine (CQ) (TCI America™).

### Bioenergetic measurements and apoptosis evaluation

ROS levels and apoptotic cell populations were quantified by flow cytometry as previously described [12]. Oxygen consumption rate (OCR) of live cells was measured in the NextGen-O2k instrument (Oroboros Instrument, Innsbruck, Austria) according to the manufacturer’s instructions. Intracellular ATP levels were assessed using CellTiter-Glo^®^ 2.0 Luminescent Cell Viability Assay (Promega). Further details on assay conditions and setup are provided in **Supplementary Methods 6**.

### Statistical analyses

Statistical analyses were performed in R or Excel. One-way ANOVA was performed in R, followed by *post hoc* Dunnett’s test, using the DMSO solvent control as the reference group. For the comparison between two groups, the homoscedastic, two-tailed Student’s *t*-test was used. All values are expressed as mean ± SEM. The number of biological replicates (n) and the statistical test applied are indicated in the respective figure legends. Significance was indicated as follows: * - *p*<0.05; ** - *p*<0.01; *** - *p*<0.001; ns : not significant (p>0.05).

## RESULTS

### *In silico* biological activity screen identifies AML-targeting compounds

Previous research in our lab identified PS127 as a novel activator of mitophagy that selectively killed AML cells [22]. A small-scale structural similarity search and structure-activity relationship analysis identified ∼15 analogs of PS127, 12 of which showed cytotoxicity against AML cells *in vitro* [12]. To identify potential mechanisms of action for these molecules, we employed the cheminformatic tool Prediction of Activity Spectra for Substances (PASS) [26]. PASS uses a training set of drug-like compounds with known biological activities to predict the activities of query molecules. PASS analysis predicted high probability for two activities, apoptotic agonism and thioredoxin reductase/glutathione reductase (T/GR) inhibition, for all 12 active PS127-series molecules [12]. Analogs inactive *in vitro* (such as PS127H) had low Pa (probability of activity) scores for these activities (**Figure 1A**). These results suggested that one or both of these activities may be involved in selective, PS127-family-mediated killing of AML cells.

**Figure 1.**
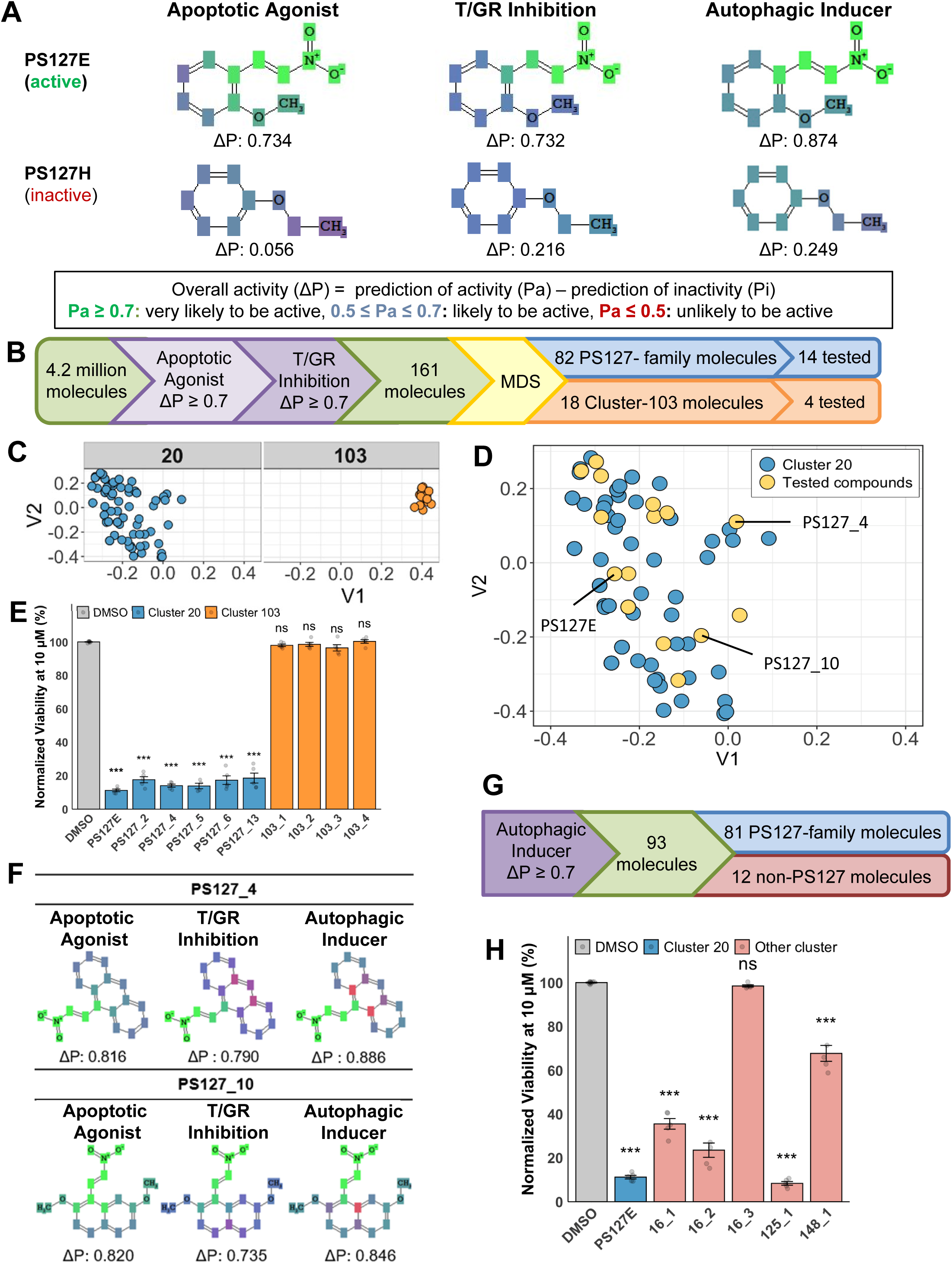
*In silico* screening and hit validation. **A)** Representative PASS outputs of active and inactive PS127 molecules on relevant biological activities. **B)** The *in silico* screening pipeline to uncover additional active novel small molecules. **C)** The top two clusters from MDS analysis. **D)** Scatter plot of cluster 20 (PS127-family), showing all PS127 molecules (blue) and tested compounds (yellow). Three molecules with the strongest cytotoxic activity are labelled. **E)** Cytotoxicity of representative molecules from cluster 20 (n=5) and cluster 103 (n=5) at 10 *μ*M on MOLM-13. **F)** PASS results of active PS127-family compounds that helped identify key third predicted activity: autophagic induction. **G)** Flow chart of a downstream *in silico* screening with the third PASS-predicted activity, autophagy inducer. **H)** Cytotoxicity comparison of compounds with three PASS-predicted activities. Representative compounds from non-PS127-family (n=5) were tested at 10 *μ*M in MOLM-13. Bar graphs display average of biological replicates, each of which is represented as data points. Error bars indicate SEM. One-way ANOVAs followed by Dunnett’s *post hoc* tests were used to assess statistical significance.

To identify additional molecules with these predicted functions, an *in silico* screen of over 4.2 million compounds was performed. To increase stringency and reduce the number of false positive hits, the screening criteria were refined. Compounds were considered hits only if ΔP (the difference between Pa and Pi, Probability of inactivity) was at least 0.7 for each of the desired activities (**Figure 1B**). This screen yielded 161 potential hits. Using structural similarity analysis and multidimensional scaling, the molecules were grouped into 41 clusters (**Supplementary Figure 1**). The majority of these compounds were within two clusters, referred to herein as cluster 20 and cluster 103 (**Figure 1C)**.

Cluster 20 contained 82 compounds, including multiple previously-identified PS127-family hits, including PS127E and PS127B. This cluster is hereafter referred to as the PS127-family. A subset of five compounds from this cluster was selected and tested as representatives of the broad chemical space of the PS127 family (**Figure 1D**). Compound cytotoxicity was tested in MOLM-13 cells at 10 µM for 72 hours (**Figure 1E**). All PS127-family compounds tested showed high cytotoxicity (**Table 1, Supplementary Table 3**).

**Table 1.** CC_50_ value of PS127-family compound in MOLM-13, PBMCs, and H9C2.

| Compound | MOLM-13 |  |  | PBMC |  |  | PBMC/MOLM-13 Ratio | H9C2 |  |  | H9C2/ |
| --- | --- | --- | --- | --- | --- | --- | --- | --- | --- | --- | --- |
| 127_1 | >8 |  |  | >16 |  |  | ND | ND |  |  |  |
| 127_2 | 1.62 | ± | 0.08 | 4.38 | ± | 0.3 | 2.71 | >16 |  |  |  |
| 127_3 | 2.41 | ± | 0.12 | 9.47 | ± | 1.56 | 3.94 | >16 |  |  |  |
| 127_4 | 0.33 | ± | 0.02 | 5.93 | ± | 0.53 | 18.04 | 5.84 | ± | 1.59 |  |
| 127_5 | 1.15 | ± | 0.05 | 7.46 | ± | 0.33 | 6.48 | >16 |  |  |  |
| 127_6 | 1.05 | ± | 0.05 | 8.37 | ± | 0.7 | 7.97 | >16 |  |  |  |
| 127_8 | 6.03 | ± | 0.22 | >16 |  |  | > 5.28 | >16 |  |  |  |
| 127_9 | 0.59 | ± | 0.02 | 9.79 | ± | 1.41 | 16.65 | 9.19 | ± | 0.88 |  |
| 127_10 | 0.63 | ± | 0.06 | 8.59 | ± | 2.42 | 13.63 | 3.73 | ± | 0.1 |  |
| 127_13 | 2.15 | ± | 0.41 | >16 |  |  | > 7.44 | >16 |  |  |  |
| 127_14 | 3.68 | ± | 0.17 | >16 |  |  | > 4.35 | >16 |  |  |  |
| 127_20 | 1.41 | ± | 0.11 | 2.94 | ± | 0.43 | 2.09 | >16 |  |  |  |
| 127_21 | >8 |  |  | >16 |  |  | ND | ND |  |  |  |
| PS127E | 0.88 | ± | 0.04 | 11.34 | ± | 1.94 | 12.89 | 16.88 | ± | 2.48 |  |

mean ± SEM, μM are shown
ND: Not determined

Cluster 103 consisted of 18 molecules, four of which were selected and tested as above (**Figure 1E**). Despite being predicted to have apoptotic agonism and T/GR inhibition, none of the compounds exhibited activity *in vitro*. Tanimoto-based structural analysis indicated that all compounds in this cluster had substantial structural similarity to each other (Tanimoto coefficient ≥0.75), whereas compounds from PS127-family cluster formed distinct sub-clusters (**Supplementary Figure 2, Supplementary Table 4**). For example, PS127_10 and PS127E were more structurally similar to each other (Tanimoto coefficient 0.833) than they were to PS127_4 or PS127_6, which were more closely related to each other (Tanimoto coefficient 0.758).

To refine our understanding of the functional differences between these clusters of compounds, PASS software was used to analyze each cluster. This process revealed several additional activities present in active, PS127-family molecules that were absent from inactive compounds (**Figure 1A and 1F**). The most intriguing of these was autophagic induction, a feature we have previously linked to mitochondrial health and AML drug sensitivity [12,27]. We hypothesized that this third activity was required for cytotoxicity and may differentiate active and inactive compounds in our group of 161 hits. This criterion excluded all the compounds in cluster 103 and reduced the number of hits to 93 (**Figure 1G**). Of these 93 molecules, 81 belonged to the PS127-family cluster, while the remaining 12 were structurally distinct from this group. Our observation that this approach removed all of Cluster 103 and narrowed our pool of hits suggested that the three activities were most directly responsible for the observed cytotoxicity. To test this hypothesis, five of the 12 non-PS127-family compounds were selected and tested for cytotoxicity in MOLM-13 cells. Four of these five significantly reduced AML cell viability **(Figure 1H)**. Three compounds, 125_1, 16_2, and 148_1, were structurally unrelated to each other and the PS127-family (**Supplementary Figure 2**), but were cytotoxic to AML cells (**Figure 1H, Supplementary Table 5**), suggesting that these compounds represent a valuable, mechanism-based toolkit for predicting cytotoxicity. These findings also provided proof of principle for the utility of using cheminformatic screening to identify compounds with desired molecular activities.

### PS127 family has selective cytotoxicity against AML cell lines

After initial cytotoxicity screening indicated that PS127-family molecules were likely to be active, 14 molecules were chosen to represent the chemical space of the PS127 family. To evaluate potency, dose-response curves were generated to determine the concentration that would induce 50% cytotoxicity (CC_50_) in MOLM-13 cells. Twelve of the 14 PS127-family molecules displayed CC_50_ under 8 µM, with four molecules (PS127_4, PS127_9, PS127_10, and PS127E) displaying sub-micromolar CC_50_ values (**Table 1, Supplementary Figure 3**).

To evaluate whether this cytotoxicity was specific to leukemic cells, CC_50_ values were also determined for healthy donor peripheral blood mononuclear cells (PBMCs). Each compound showed at least two-fold higher CC_50_ values in PBMCs compared to MOLM-13. Notably, CC_50_ values of lead PS127-family compounds in PBMCs were at least 12- and 8-fold higher than in MOLM-13 and MV4;11, respectively, indicating consistent compound potency across AML cell lines harboring FLT3-ITD mutations.

Previous research indicated that PS127 (the founding member of the compound family)likely functions by stimulating autophagic recycling of mitochondria [12]. We considered the possibility that these compounds may exhibit greater toxicity to cells with increased mitochondrial dependence, such as cardiomyocytes [28]. To test this, CC_50_ values were also calculated for H9C2 cardiomyocytes for PS127-family compounds with sub-micromolar CC_50_ values in MOLM-13 cells. Observed CC_50_ values were ∼5 to 19-fold higher for cardiomyocytes than AML cells, suggesting that cytotoxicity was specific to leukemic cells due to their unique metabolic adaptations **(Table 1)**.

These compounds were also evaluated in a panel of other AML cell lines, including MOLM-14, MV4;11, NB4, and OCI-AML2 (**Table 2**), which have a variety of genetic lesions frequently found in AML patients. Each of the prioritized PS127-molecules was active against other AML cell lines, a result crucial for targeting the heterogeneous nature of AML. Notably, PS127E and PS127_10 displayed similar activity in large-scale dose response analysis in MOLM-13 and MV4;11 and much lower toxicity to PBMCs (**Supplementary Figure 3).** Based on their strong activity in AML cell lines, PS127_4, PS127_10, and PS127E were selected for further evaluation.

**Table 2.** CC_50_ value of prioritized PS127-family compounds in other AML cell lines.

| Compound | MOLM-14 |  |  | MV4;11 |  |  | NB4 |  |  | OCI-AML2 |  |  |
| --- | --- | --- | --- | --- | --- | --- | --- | --- | --- | --- | --- | --- |
| **127_4** | 0.71 | ± | 0.04 | 0.73 | ± | 0.07 | 0.9 | ± | 0.07 | 0.96 | ± | 0.14 |
| **127_9** | 0.94 | ± | 0.09 | 1.38 | ± | 0.12 | 1.36 | ± | 0.06 | 1.38 | ± | 0.11 |
| **127_10** | 0.92 | ± | 0.05 | 0.63 | ± | 0.03 | 0.93 | ± | 0.04 | 1.11 | ± | 0.11 |
| **PS127E** | 1.28 | ± | 0.09 | 1.42 | ± | 0.01 | 1.23 | ± | 0.14 | 2.35 | ± | 0.23 |

mean ± SEM, μM are shown

### PS127-family compounds are synergistic with current commercial chemotherapeutics

Due to the prevalence of drug resistance in AML, most modern chemotherapeutics are administered in combinations. The synergy of our top compounds (greater-than-additive effect) was tested in combination with commonly-used AML chemotherapeutics [29], including the first-line treatments doxorubicin (DOX), cytarabine (ara-C), venetoclax (VEN) [30–32], and midostaurin (MID), a targeted therapy used in AML patients with FLT3 mutations [33,34]. Synergy was evaluated by assessing cell viability after single and combinatorial treatments using a Bliss independence model, with a score above 10 considered to indicate synergy [29]. The prioritized candidates, PS127_4, PS127_10, and PS127E, showed some synergy with DOX and ara-C but had the strongest synergy with MID in MOLM-13 (**Figure 2A-D, Supplementary Figure 4)**.

**Figure 2.**
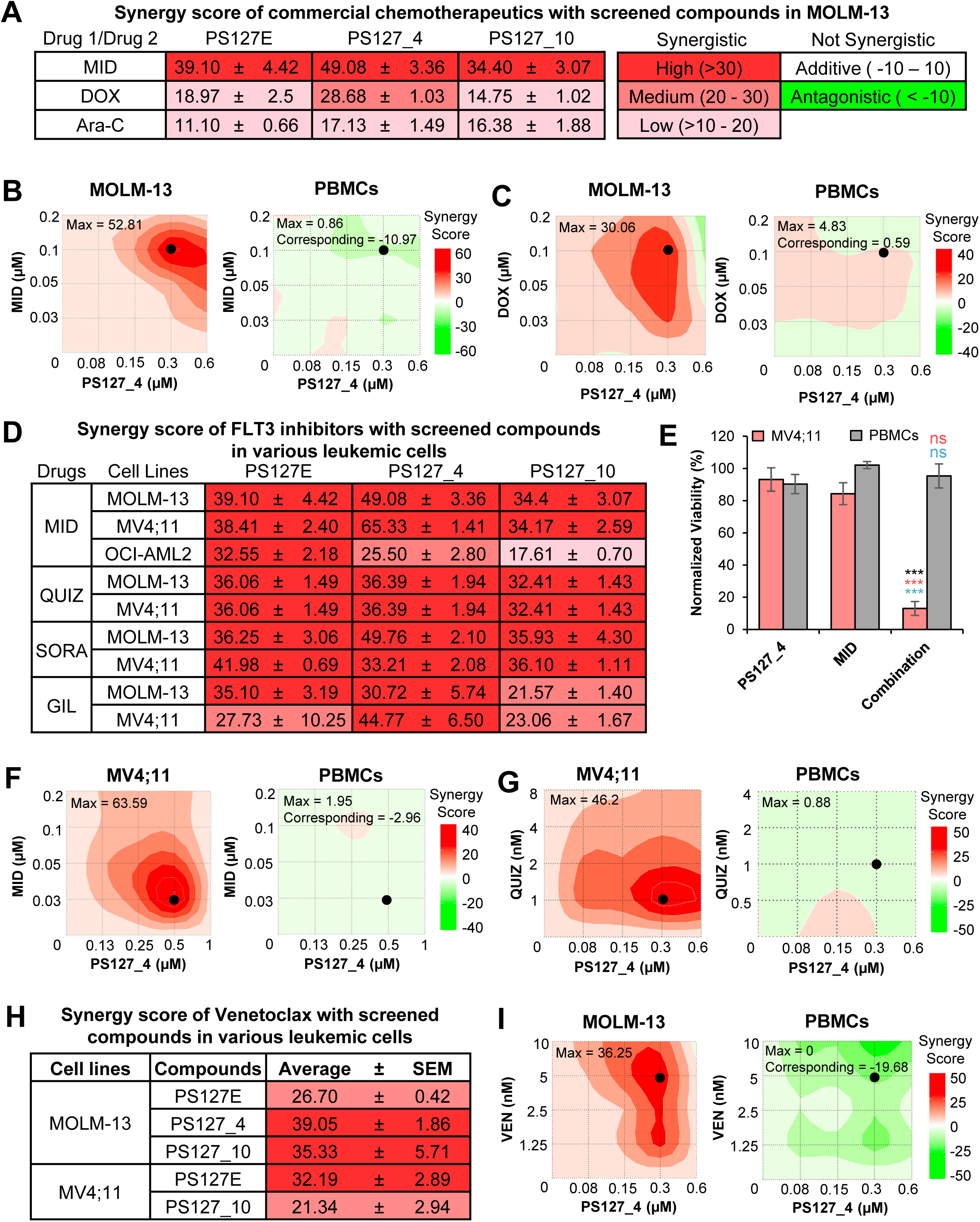
Synergistic effect of PS127-compounds and commercial chemotherapeutics. **A**) Average maximum synergy score (± SEM) of PS127-family compounds and commercial chemotherapeutics: MID, DOX, and ara-C (n = 3). **B**) Representative synergy plot of PS127_4 and MID in MOLM-13 cells (left) or PBMCs (right). **C**) Representative synergy plot of PS127_4 and DOX in MOLM-13 cells (left) or PBMCs (right). **D**) Average maximum synergy score (± SEM) of PS127-family compounds and various FLT3 inhibitors (n = 3). **E**) Comparison of MV4;11 and PBMCs viability upon monotherapy or combined treatments with PS127_4 and MID. All bar graphs represent mean ± SEM from three biological replicates (n = 3). Black asterisks indicate comparison of AML cells vs. healthy PBMCs under the same combinatorial treatment conditions. Pink asterisks indicate significantly lower survival under combinatorial treatment compared to single PS127-family compound treatment; Blue asterisks indicate significantly lower survival under combinatorial treatment compared to a single MID treatment. Black dot represents maximum synergy point in leukemic cells and its corresponding condition in PBMCs. Significance was assessed via Student’s *t*-test. **F**) Representative synergy plot of PS127_4 with MID in MV4;11 (left) and PBMCs (right). **G**) Representative synergy plot of PS127_4 with QUIZ in MV4;11 (left) and PBMCs (right). **H**) Average maximum synergy score (± SEM) of PS127-family compounds with VEN in AML cells (n = 3). **I**) Representative synergy plot of PS127_4 and VEN in MOLM-13 cells (left) or PBMCs (right).

We also evaluated the synergistic potential of these compounds in healthy PBMCs under the same conditions. In these cells, no synergistic activity was observed (**Figure 2B-C, Supplementary Figure 4)**. We found that the survival in the presence of drug combinations resulting in maximum synergy was significantly lower in AML cells than in PBMCs, with survival in the latter remaining >80% for these tested combinations **(Figure 2E, Supplementary Figure 5**). This reinforced that drug combination cytotoxicity was specific to leukemic cells and is unlikely to have overt side effects on healthy cells.

Since PS127-family compounds showed the greatest synergy with MID, we evaluated these combinations in two additional AML cell lines: MV4;11 and OCI-AML2. MV4;11, like MOLM-13, carries an FLT3-ITD mutation, while OCI-AML2 lacks known FLT3 mutations and maintains detectable amounts of FLT3 [35]. As anticipated, top PS127-compounds showed significant synergy with MID in MV4;11 (**Figure 2F, Supplementary Figure 6**). The combination of PS127_4 and MID produced the strongest synergy, with the average maximum Bliss score being above 60 (**Figure 2D,F**). This combination did not significantly affect PBMC cell viability (**Figure 2E**). The difference in combined PS127_4 and MID treatment survival between MV4;11 and PBMCs was approximately 7-fold (13% vs 95%). Notably, prioritized PS127-compounds displayed a range of synergy with MID in OCI-AML2, despite the lack of FLT3 mutation in this cell line (**Figure 2D, Supplementary Figure 6C-E**). MID is also known to target S6 phosphorylation and sphingosine kinase-1, which may contribute to its interaction with PS127-family compounds in OCI-AML2 cells [35,36]. Importantly, while PS127E had comparable synergy with MID in MV4;11 and OCI-AML2, synergy scores for PS127_4 and PS127_10 were approximately two-fold lower in the latter cell line, indicating complex relationships between compound effects and cancer genotype.

Due to the strong synergy with MID, we evaluated the synergistic potential of PS127E, PS127_4, and PS127_10 with other FLT3 inhibitors, quizartinib (QUIZ), sorafenib (SORA), and gilteritinib (GIL). We found that these compounds had strong synergy with QUIZ and SORA **(Figure 2D,G, Supplementary Figure 7A-H)** and ranged from moderate to strong synergy with GIL **(Figure 2D, Supplementary Figure 7I-L)** in both AML cell lines. These data indicate that our PS127-family compounds will likely be active in AML patient cells, regardless of whether they carry wild-type or mutant FLT3 alleles.

Furthermore, we evaluated the synergistic potential of lead compounds in combination with venetoclax VEN, an established standard-of-care agent for newly-diagnosed and refractory/relapsed AML [37]. All leads demonstrated strong synergy in MOLM-13 cells (**Figure 2H-I, Supplementary Figure 8**). Importantly, PS127_4 showed the strongest synergy with VEN in leukemic cells while remaining antagonistic in PBMCs (**Figure 2I**). Similar synergistic activity was observed in FLT3-ITD MV4;11 cells. These data are consistent with a dual-action mechanism wherein PS127-mediated cellular stress sensitizes cells to VEN-induced apoptosis, highlighting the potential for these combinations in broader clinical applications.

### *In vitro* validation of PASS activity predictions

To elucidate the mechanisms used by the prioritized compounds to selectively induce leukemic cell death, we first verified the PASS-predicted activities used for screening: apoptotic agonism, T/GR inhibition, and autophagic induction.

To evaluate compounds’ ability to induce apoptosis, flow cytometry experiments were performed using annexin V-FITC/ Hoechst/PI staining. Flow cytometry analysis demonstrated that 24-hours treatment with PS127E, PS127_4, or PS127_10 significantly induced apoptosis in leukemic cells, as indicated by annexin V-positive staining (**Figure 3A-B**), suggesting that apoptosis is a primary mechanism of cell death.

**Figure 3.**
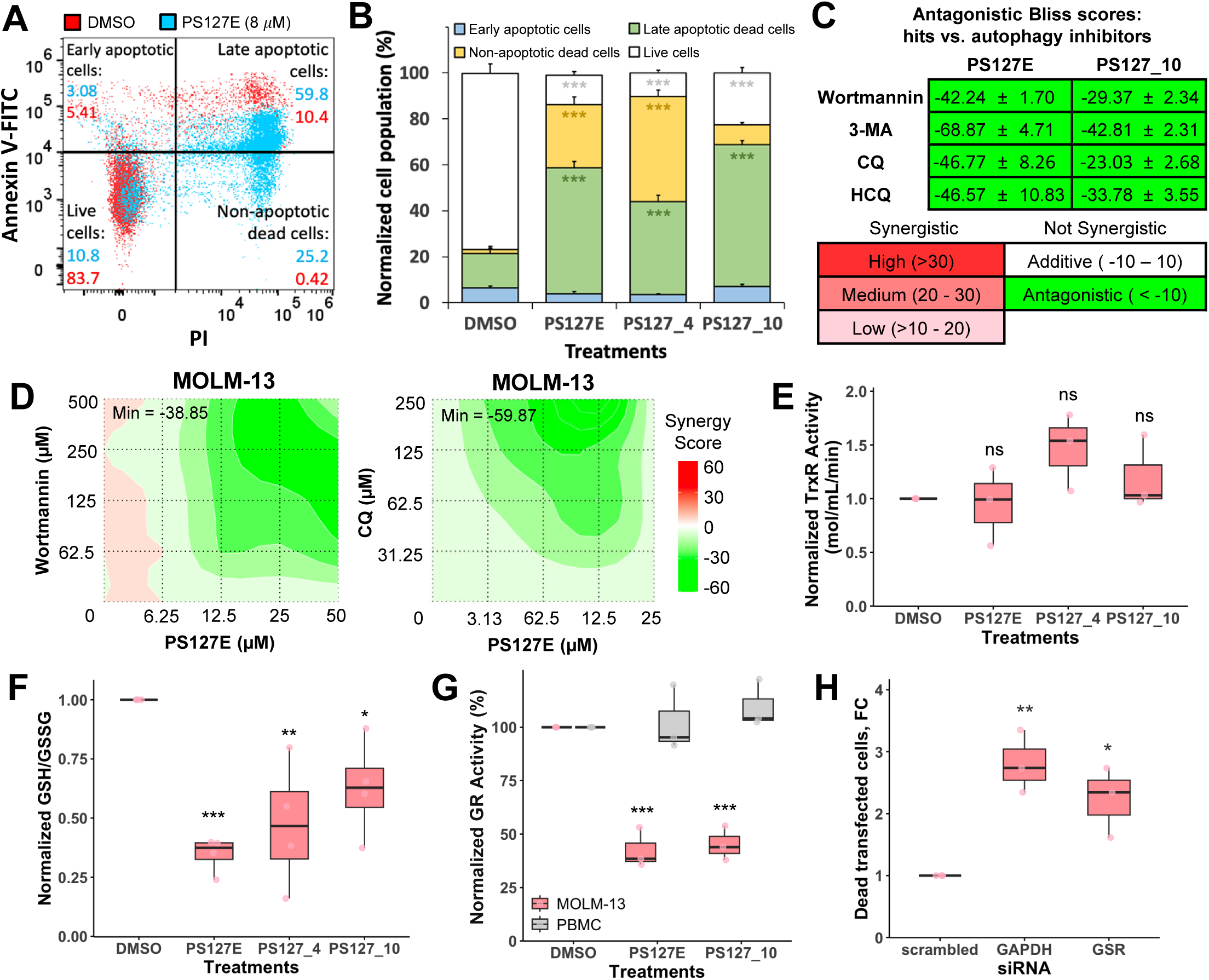
Validating predicted activities of PS127-treatment in MOLM-13. **A**) Representative flow cytometry analysis of apoptotic cells induced by PS127E (8 *μ*M, blue) or DMSO (red) for 24 hours. Apoptotic assay was determined using flow cytometry with Annexin V-FITC/PI/Hoechst staining. **B**) Quantification of cell populations upon PS127-molecule treatment compared to DMSO following annexin V-FITC/PI staining (n = 4). Apoptotic cells were detected by annexin V-FITC^+^/PI^-^ and annexin V-FITC^+^/PI^+^, while non-apoptotic dead cells were indicated by annexin V-FITC^-^/PI^+^ and live cells were annexin V-FITC^-^/PI^-^. Bar graphs show the average percentage of cell population and error bars display SEM. Significance levels represent comparison of PS127-induced apoptotic cells to DMSO. **C**) Average antagonist score (± SEM) of PS127-molecules with various autophagic inhibitors (n = 3). **D)** Representative antagonism landscape of PS127E and wortmannin (left) or CQ (right) treatment. **E**) The effect of PS127-moleculeson TrxR activity (n = 3). **F**) Changes of GSH to GSSG ratio upon treatment of PS127-molecules (n= 4). **G**) Quantification of GR enzymatic activity upon PS127-molecules treatment in MOLM-13 and PBMCs. **H**) Fold change of dead transfected MOLM-13 cells after *GSR* knockdown relative to scrambled siRNA. Biological replicates are shown as data points. One-way ANOVAs followed by Dunnett’s *post hoc* tests were used to assess statistical significance.

Next, the dependence of PS127-family compound activity on autophagic stimulation was evaluated. MOLM-13 cells were treated for 24 hours with PS127-family molecules and either early-stage (wortmannin) and 3-methyladenine [3-MA]) or late-stage (chloroquine [CQ] and hydroxychloroquine [HCQ]) inhibitors of autophagy [38–40]. The addition of autophagic inhibitors to PS127-family-treated cells resulted in strong antagonism, suggesting that functional autophagy was required for PS127-induced cytotoxicity (**Figure 3C-D, Supplementary Figure 9**).

Finally, we evaluated the predicted inhibitory effect of PS127-family molecules on enzymes responsible for redox metabolism, specifically TrxR or GR. None of the prioritized PS127-family compounds had a significant effect on the activity of TrxR in MOLM-13 (**Figure 3E**), suggesting that none of them interacted with TrxR.

In contrast, compound-mediated effects on glutathione metabolism were observed. The GSH pool maintains ROS homeostasis in the cell and reduces oxidative stress [41,42]. GSH is oxidized to GSSG, which is then reduced back to GSH by GR, restoring its potential to serve as an antioxidant. The steady-state ratio of GSH to GSSG represents the current oxidative burden of the cells, but can also serve as a readout of GR activity. Cells treated with PS127-family compounds exhibited significantly decreased GSH-to-GSSG ratios (**Figure 3F**), suggesting that the compounds may be disrupting the function of the enzyme.

To test the specificity of this activity *in vitro*, MOLM-13 or PBMC cells were treated with PS127-family compounds for 24 hours and then lysed. GR activity in cell-free lysates was quantified and normalized to total protein concentration, in accordance with GR activity assay. Lead compounds significantly inhibited GR activity in leukemic cells, but not in PBMCs (**Figure 3G**), likely reflecting increased basal GR activity in leukemic cells. By inhibiting GR activity, lead compounds prevented recycling GSSG to GSH thereby increasing oxidative stress and cytotoxicity.

To confirm the importance of GR for survival in MOLM-13 cells, siRNAs were used to knock down *GSR* or *GAPDH* (the latter served as a positive control). Disruption of *GSR* resulted in considerable cytotoxicity in MOLM-13 cells compared to scrambled siRNA (**Figure 3H**). This corroborates our observations with chemical inhibition and reinforcing the conclusion that GR activity is critical for survival of MOLM-13 cells. Unfortunately, a limited number of viable cells were obtained after *GSR* knockdown, precluded determination of CC_50_ values in *GSR* knockdown cells.

### PS127-compounds induce redox-dependent cytotoxicity by interfering with glutathione metabolism

To further validate the mechanism of compound activity on redox metabolism, PS127E-treated cells were supplemented with exogenous GSH, GSSG, or the GSH precursor, *N*-acetyl-cysteine (NAC). GSH and NAC completely rescued viability of PS127E-treated cells (**Figure 4A**). In contrast, supplementation with GSSG did not provide any rescue. As neither GSH nor GSSG are toxic to MOLM-13 (**Supplementary Figure 10A**), we hypothesized that this difference is directly related to their distinct functions in redox metabolism.

**Figure 4.**
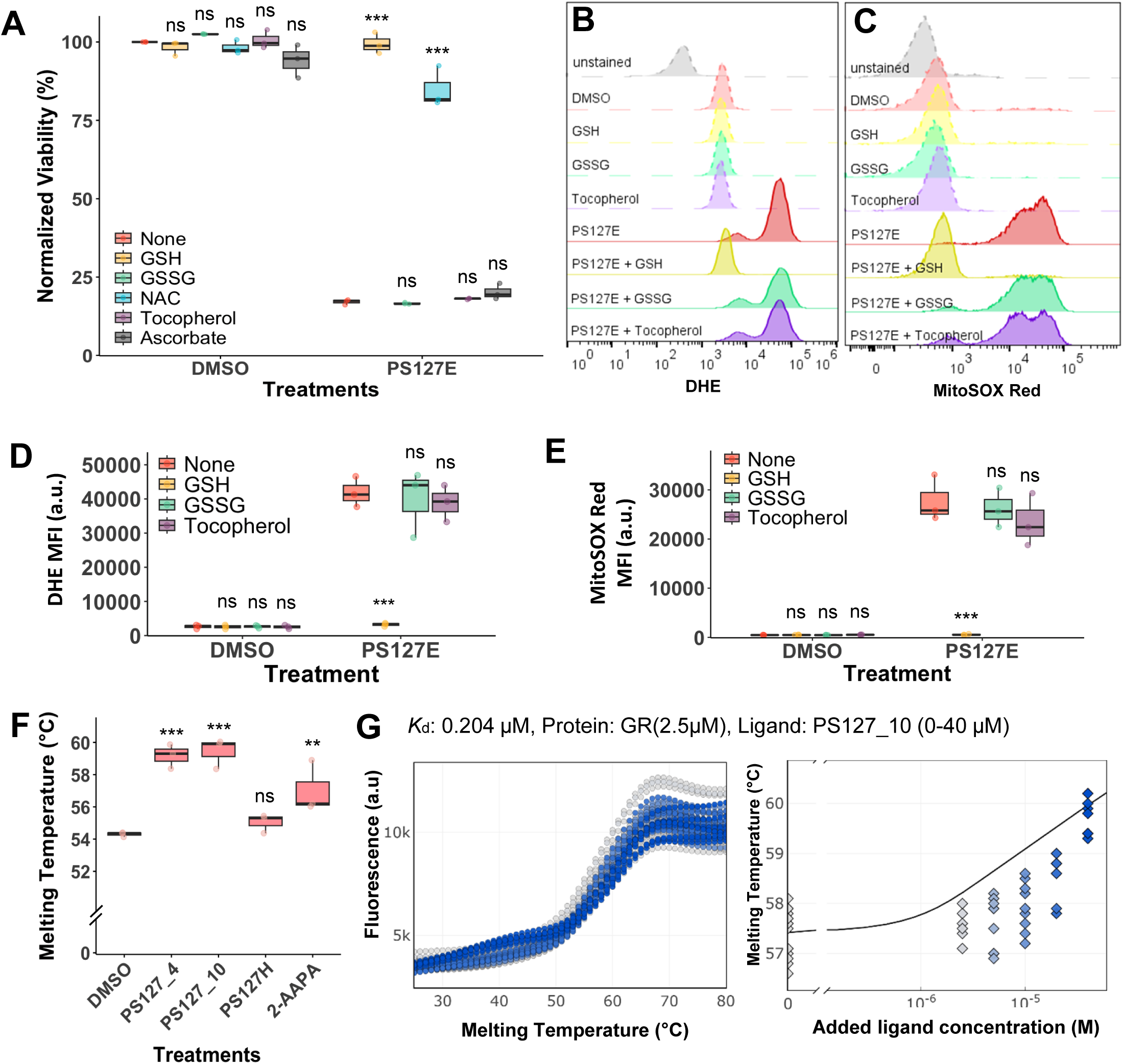
The effect of PS127-induced cytotoxicity on redox metabolism and ROS levels. **A**) Redox metabolite-mediated rescue of cell viability upon PS127E treatment (n = 3) for 72 hours. **B**) A representative histogram of DHE^+^ population in the presence of redox metabolites. **C)** Representative histogram of MitoSOX Red^+^ population in the presence of redox metabolites. Peaks with dashed lines indicate control treatment, while solid lines represent treatment with PS127-molecule. **D**) Total ROS quantification using median fluorescence intensity (MFI) of DHE in the presence of redox metabolites in the presence and absence of PS127E (n = 3). **E**) Mitochondrial ROS quantification using MFI of MitoSOX Red in the presence of redox metabolites and presence and absence of PS127E (n = 3). ROS level was indicated by MFI of ROS staining dye. **F**) Melting temperature shift of human GR in the presence of PS127-family compounds. PS127_4 (n = 4), PS127_9 (n = 3), PS127_10 (n = 3), and 2-AAPA (n = 3) significantly changed the melting temperature of GR compared to DMSO (n = 7), indicating compound-enzyme binding. PS127H (n = 3) did not change the melting temperature. PS127H = previously identified inactive PS127 analog, 2-AAPA = GR inhibitor control. **G)** Raw fluorescence of Sypro Orange for technical replicates of the DSF to determine K_d_ between PS127_10 and GR (left). Melting temperature of GR with PS127_10 at 0, 2.5, 5, 10, 20, and 40 µM (right). One-way ANOVAs followed by Dunnett’s *post hoc* tests were used to assess statistical significance.

To verify that the increase in viability conferred by GSH or NAC was not simply a consequence of providing antioxidants to MOLM-13 cells, MOLM-13 cells were treated with PS127-family compounds and then supplemented with other known antioxidants that act independently of GR. Reduced ascorbate, which does not affect glutathione peroxidase or GR activity [41,43], was tested at the highest concentration that could be used without causing significant cytotoxicity on its own (**Supplementary Figure 10B**). Even at this concentration, the ascorbate did not significantly affect cell death (**Figure 4A**). Supplementation with α-tocopherol, another antioxidant [44,45], also failed to significantly improve survival of PS127E-treated cells (**Supplementary Figure 10C, Supplementary Figure 11A-B**). The combination of ascorbate and α-tocopherol generates enhanced ROS scavenging activity [46]. Interestingly, this combination still did not rescue cells from PS127-induced cytotoxicity (**Supplementary Figure 11B**), Likewise, the cytotoxicity of PS127_4 and PS127_10 was inhibited by GSH but remained unaffected by GSSG or α-tocopherol, ascorbate, or their combination (**Supplementary Figure 11A-B**). These results reinforced the likelihood that PS127-family molecules specifically compromise the glutathione pathway, likely by preventing GR from reducing GSSG, and that this effect cannot be compensated for by other antioxidative pathways.

GSH serves as a scavenger for intracellular and mitochondrial ROS, therefore its depletion would be expected to increase both total and mitochondrial ROS levels [47]. To examine the relationship between ROS levels and cytotoxicity, total and mitochondrial ROS were measured in cells treated with PS127-family molecules using dihydroethidium DHE and MitoSOX Red staining, respectively. PS127-family compound treatment increased both total and mitochondrial ROS levels (**Figure 4B-E, Supplementary Figure 11C-H**). Consistent with the viability rescue data, only supplementation with GSH, but not with GSSG or α-tocopherol, restored total and mitochondrial ROS levels in PS127-family-treated cells to approximately basal levels.

### PS127-family compounds interfere with glutathione reductase

This evidence strongly suggests that GR is the target of the PS127-family compounds. To directly prove physical interaction between the compounds and GR, we used differential scanning fluorimetry (DSF). This technique measures the melting temperature of a target protein in the presence or absence of small molecules that may interact with the protein [48]. Changes to the melting temperature of the protein indicate stabilization of the protein, likely due to interaction between the enzyme and the small molecule.

Incubation of GR with active PS127-family compounds (PS127_4, PS127_10, or PS127E) significantly increased its melting temperature, suggesting that the compounds interact with the enzyme (**Figure 4F**). In contrast, PS127H, an inactive PS127-like compound, did not significantly alter the melting temperature of GR, indicating that the lack of activity is likely due to a lack of interaction between GR and PS127H. Consistent with these findings, 2-AAPA, a known inhibitor of GR [49], produced a comparable increase in melting temperature.

To further verify this interaction, a thermal shift binding assay was performed between PS127_10 and purified GR, which demonstrated a concentration-dependent increase in melting temperature **(Figure 4G)**. Using these data, a binding constant (*K_d_*) for the experimental conditions was calculated to be 0.204 µM [95% CI: 0.160-0.247 µM], indicating high-affinity binding of PS127_10 to GR under the experimental conditions used (2.5 µM GR). These data indicate that active PS127-family molecules bind to GR, inhibiting GSSG reduction, resulting in depletion of cellular GSH pools.

### A non-PS127-family compound, 125_1, shares PS127-family compounds’ properties

As described above, the PS127 hits tested were predicted to, and were confirmed to, have high probability of apoptotic agonism, T/GR inhibition, and autophagic induction. These effects are likely to have driven the observed cytotoxicity to MOLM cells. The screen that identified several of these hits was structure-agnostic, so other compounds with all three activities may also be cytotoxic.

To test this prediction, we procured compound 125_1, which is structurally unrelated to the PS127-family (Tanimoto coefficient < 0.7, **Supplementary Table 4**) but was predicted to have all three activities. Compound 125_1 exhibited a low CC_50_, 0.41 ± 0.04 µM in MOLM-13, which was over 30-fold lower than its CC_50_ for PBMCs (15.68 ± 0.75 µM) or cardiomyocytes (>16 µM). Like PS127-family compounds, 125_1 induced apoptotic cell death, was antagonistic with 3-MA, its cytotoxicity was inhibited by GSH, but not by GSSG or α-tocopherol (**Figure 5A-C**). Compound 125_1 also increased total and mitochondrial ROS levels, an effect attenuated by GSH (**Figure 5D-G**). Finally, compound 125_1 reduced GR activity (**Figure 5H**), likely by directly binding to GR, as shown by the protein’s increased melting temperature in thermal shift assay (**Figure 5I**). Compound 125_1 also showed synergy with MID in MOLM-13, like PS127-family molecules (**Figure 5J**). Overall, these data indicate that 125_1 and the PS127 family share the same mechanistic properties, despite structural dissimilarity.

**Figure 5.**
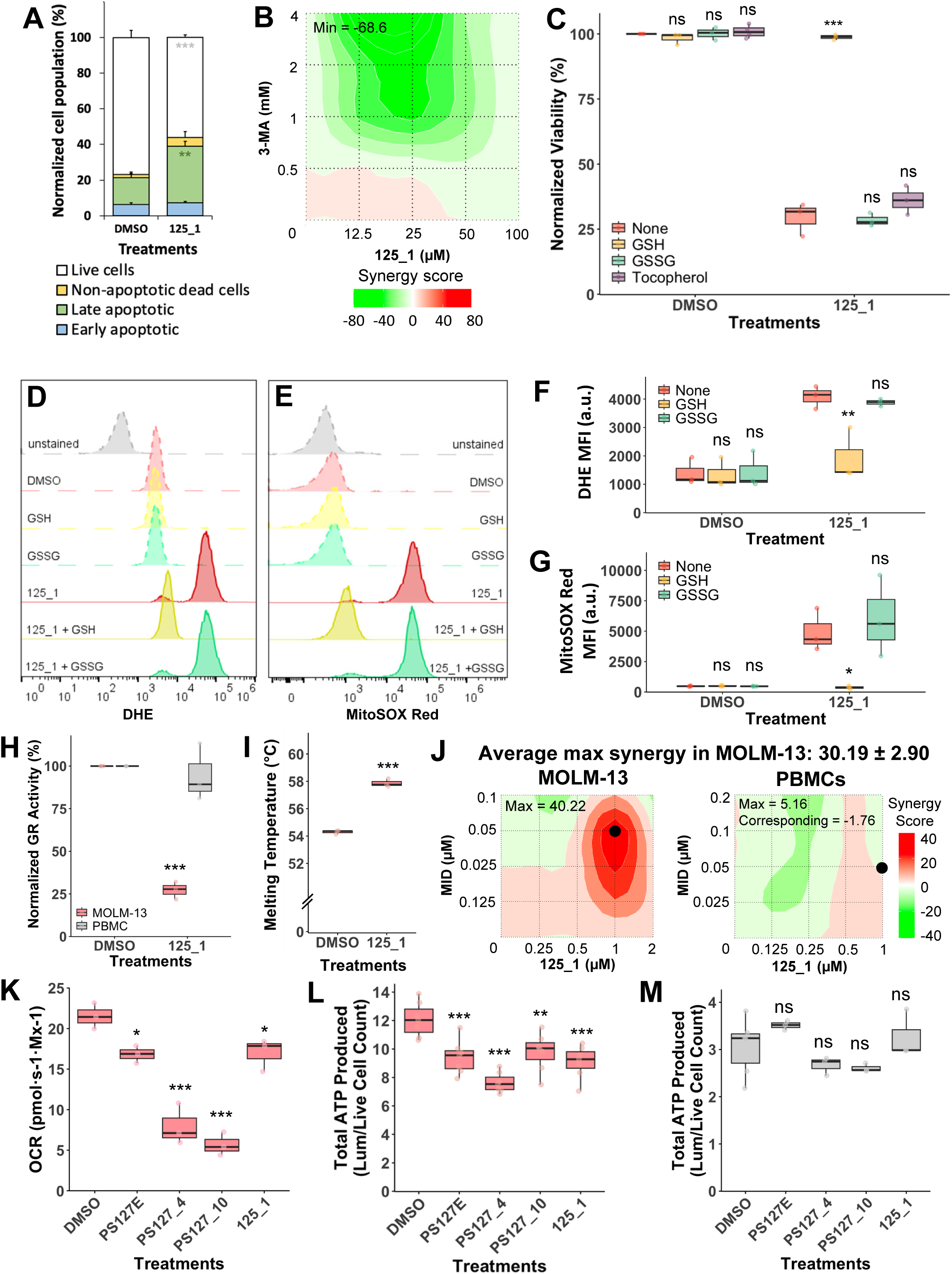
Evaluation of PASS-predicted activities in 125_1 and the impact of PS127-family compounds and 125_1 on mitochondrial bioenergetics. **A**) Quantification of apoptotic cells upon 125_1 treatment in MOLM-13, indicated by Annexin V/PI staining (n = 4). **B**) Representative antagonistic interactions between 3-MA and 125_1. **C**) Cytotoxicity upon 125_1 treatment was rescued by GSH (n = 8). **D**) DHE^+^ population representative histogram upon 125_1 treatment in the presence or absence of redox metabolites. **E)** MitoSOX Red^+^ population representative histogram upon 125_1 treatment in the presence or in the absence of redox metabolites. Peaks with dashed lines indicate control treatment or in absence of PS127-molecule, while solid lines represent treatment with PS127-molecule. **F**) Quantification of total ROS in the presence of redox metabolites (n = 3) based on MFI of DHE. **G**) Quantification of MitoSOX Red MFI in the presence of redox metabolites (n = 3). **H**) Quantification of GR activity in MOLM-13 treated with 125_1 (n = 3). **I**) Melting temperature shift of human GR upon 125_1 treatment in MOLM-13 (n = 5). **J**) Representative synergy plot of compound 125_1 and MID in MOLM-13 (left) and PBMCs (right). **K**) OCR measurement of MOLM-13 upon 3-hour compound treatment. PS127E (n = 4), PS127_4 (n = 5), PS127_10 (n = 4), or 125_1 (n = 3) impaired the routine OCR compared to DMSO (n = 7). OCR was normalized to the number of live million cells per chamber volume. **L**) The steady state ATP level in MOLM-13 after 3-hour exposure of compound (n = **7**). **M**) The steady state ATP level in PBMCs after 3-hour exposure of compound (n = 3). All AML cells and PBMCs had > 90% viability before cell lysis occurred to measure total ATP. One-way ANOVAs followed by Dunnett’s *post hoc* tests were used to assess statistical significance of apoptotic cells, ROS and ATP levels, and oxygen consumption. Significance level of DSF was determined using Student’s *t*-test.

### Prioritized PS127-family compounds interfere with mitochondrial bioenergetics

As noted above, PS127-family compounds compromised redox homeostasis and increased mitochondrial ROS, suggesting that the compounds may be affecting mitochondrial function, particularly respiration. AML cells, especially LSCs, have low coupling efficiency and therefore rely heavily on OXPHOS to support their increased demand for ATP [12,50,51]. We previously reported that other compounds that impair respiration and ATP production exhibit selective cytotoxicity [12].

To test whether PS127-family compounds similarly impaired mitochondrial respiration, oxygen consumption was measured in MOLM-13 cells that were exposed to PS127_4, PS127_10, PS127E, or 125_1 for 3 hours. Each of the compounds impaired routine respiration, as shown by reduced OCR (**Figure 5K**). Consistent with previous results [12], all lead compounds significantly reduced ATP production in MOLM-13, but not in PBMCs (**Figure 5L-M**).

### PS127-family activity in AML patient samples

Patient-derived samples were used to validate PS127-family compound activity in a more clinical context. In a panel of wild-type and FLT3-mutant samples, we evaluated the ability of compounds to induce cytotoxicity, synergy between PS127_10 and MID or DOX, oxygen consumption rate, and rescue by exogenous GSH. Due to limited cell numbers in patient-derived samples, limited experiments could be performed.

In line with the heterogeneity of AML in patients, the OCR of the primary samples ranged from ∼5 to ∼30 pmol/s per million viable cells; MOLM-13 and PBMC OCR values were∼18 and ∼3 pmol/s per million viable cells, respectively (**Figure 6A**, **6F**). Additionally, GSH supplementation rescued AML patient cells treated with PS127E or compound 125_1, regardless of FLT3 mutation status (**Figure 6B-C**). A range of effects, from additivity to strong synergy, were seen in various patient-derived cells when PS127_10 was combined with MID or DOX (**Figure 6D-F**). FLT3 mutation status did not appear to correlate with the degree of synergy.

**Figure 6.**
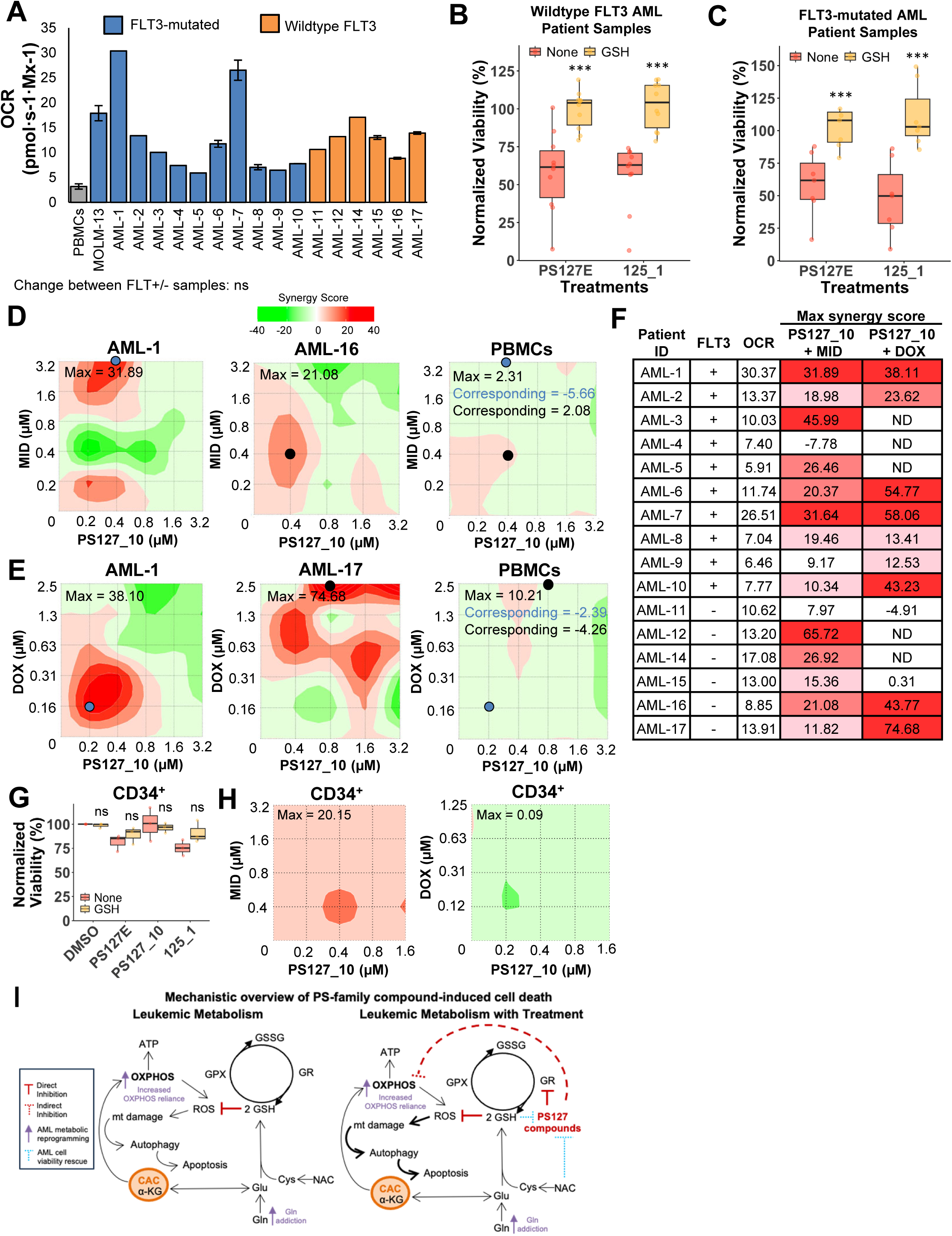
Screen hits are effective in AML patient samples. **A)** OCR of FLT3-mutated and wild-type AML patient cell samples, PBMCs, and MOLM-13 in the 0.5 mL chamber. OCR was normalized to the number of live million cells per mL [pmol·s^-1^ per million cells (Mx^-1^)]. Error bars represent SEM from two technical replicates. Bars without error bars represent one replicate due to limited sample size. **B)** Viability rescues of PS127E- or 125_1-treated wild-type AML patient cells (n = 10) by 500 µM GSH supplementation. **C)** Viability rescues of PS127E- and 125_1-treated FLT3-mutated patient cells (n = 7) by 500 µM GSH supplementation. **D**) Representative synergy plot of FLT3-mutated (AML-1) (left), wild-type FLT3 (AML-16) (middle), and PBMCs (right) of PS127_10 and MID.**E**) Representative synergy plot of FLT3-mutated (AML-1) (left), wild-type FLT3 (AML-17) (middle), and PBMCs (right) of PS127_10 and DOX. Blue and black dots represent the maximum FLT3-mutated synergy point and its corresponding condition in PBMCs. Black dots correspond to maximum wild-type FLT3 synergy point and its corresponding PBMC condition. **F**) Summary table of OCR and max synergy value with PS127_10 and MID or DOX in AML patient cells. ND = not determined. **G**) The effect of 500 µM GSH supplementation on CD34^+^ cells treated with lead compounds (n = 3). **H**) Representative synergy plot of PS127_10 and MID in CD34^+^ cells. **I**) Hypothesized mechanistic model of active PS127-family compounds inhibiting GR, causing an increase in ROS, leading to lethal mitophagy. Leukemic metabolism without (left) and with (right) PS127-compound treatment are shown. ATP = adenosine-triphosphate, OXPHOS = oxidative phosphorylation, GSH = reduced glutathione, GSSG = oxidized glutathione, NAC = N-acetylcysteine, Gln = glutamine, Glu = glutamate, Cys = cysteine, CAC = citric acid cycle, α-KG = α-Ketoglutarate, GPX = glutathione peroxidase.

### Activity of lead compounds in human bone marrow-derived CD34^+^ progenitors

To further evaluate the therapeutic potential of these compounds, lead compounds were tested on human bone marrow-derived CD34^+^ cells, which are related to the CD34^+^CD38^-^ population that gives rise to LSCs [52–54], enabling evaluation of potential off-target effects on healthy hematopoietic progenitors. PS127E and PS127_10 exhibited significantly higher median CC_50_ values, 14.73 and 39.15 μM, respectively, than were seen for AML cells. Unlike MOLM-13 cells, GSH supplementation had negligible effect on the viability of the CD34^+^ HSCs (**Figure 6G**), likely due to the low cytotoxic effect of compound, indicating that the redox status of these cells has not been compromised. PS127_10 showed a moderate synergistic effect with MID and no synergy with DOX in CD34^+^ progenitor cells **(Figure 6H)**, markedly weaker than the synergistic interaction observed in MOLM-13 cells, suggesting that healthy cells lack the specific redox and signaling susceptibilities that are exploited in drug cooperation and further supporting the conclusion that treatment with these compounds will specifically target cancerous cells with minimal impact on HSCs.

## DISCUSSION

It remains imperative to identify new, improved treatments for AML that can be more broadly tolerated by patients who are elderly or have comorbidities that preclude standard induction and consolidation. To more efficiently probe chemical space and identify promising leads, researchers leverage *in silico* techniques like ligand- and structure-based screening methods [55]. Structure-based screening is popular, but is limited to targets where comprehensive structural information is available [56]. In its absence, researchers are forced to rely on predicted structures gained from homology modeling or template-free, AI-based techniques like AlphaFold. Ligand-based approaches such as quantitative structure-activity relationship (QSAR) modeling or molecular docking are also common, but suffer from similar limitations [57,58].

To circumvent this limitation, function-based methods, like PASS, have been developed. PASS is a ligand-based cheminformatic approach trained on structural-activity relationships for thousands of known bioactive compounds, enabling rapid prediction of the biological functions for millions of as yet uncharacterized molecules [59,60]. This method can facilitate a more approachable cheminformatic study with a target-agnostic approach, conceptually similar to the high-throughput, high-content phenotypic screening we have previously performed in other contexts [22,61].

This approach was effective both for predicting biochemical activities for our hit compounds and for identifying ∼160 hits in an *in silico* screen. Moreover, counter-analysis of inactive compounds identified a third activity (autophagic induction) that ruled out almost half of the initial hits, reducing time and money spent on laborious and expensive follow-up studies.

Mitochondria represent promising therapeutic targets for leukemia treatment due to the metabolic programming observed in AML cells, which includes increased reliance on OXPHOS and glutaminolysis over glycolysis. These characteristics render the mitochondria in AML cells partially dysfunctional and susceptible to damage (63). Mitochondrial damage often leads to mitophagy, the selective autophagic recycling of damaged mitochondria to replenish cellular metabolite pools for use as building blocks (49). This process can help maintain cellular health or be lethal (103). In line with this observation, mitophagic activators and inhibitors can both induce selective leukemia cell death. Additionally, current chemotherapeutics also induce mitophagy indirectly (114).

Accordingly, targeting redox homeostasis has become an increasingly appealing approach to treat heterogeneous cancers like AML. Interestingly, our lead compounds showed synergy with various FLT3 inhibitors, which is likely due to dual activity targeting the cellular redox homeostasis and FLT3 signaling. We propose that GR inhibition acts as a metabolic sensitizer. Regardless of their FLT3 genotype, we observed increased OCR in AML cells compared to PBMCs (**Supplementary Figure 12**). By concurrently inhibiting GR (**Figures 3G**, **5H**), lead compounds reduce GSH pool that helps to mitigate oxidative stress. Consequently, this sensitizes cells to mitochondrial disruption caused by FLT3 inhibition. Furthermore, the FDA-approved BCL-2 inhibitor venetoclax (VEN, ABT-199), is a first-in-class AML chemotherapeutic [37,62,63]. BCL-2 regulates mitochondrial respiration; accordingly, inhibitors like VEN, ABT-737, and ABT-263 selectively induce cell death in leukemic cells with low ROS while having markedly less effect on PBMCs [6]. Other studies have identified compounds that induce leukemic cell death by interfering with glutathione metabolism, specifically by interfering with glutaminase, glutamyl cysteine synthetase, and glutathione peroxidase [10,64,65]. Our compounds produce similar phenotypic effects, like decreased glutathione levels and elevated ROS levels through the unique mechanism of GR inhibition. Previously, we showed that PS127B and PS127E activate high levels of apoptosis in LSCs but not in healthy hematopoietic stem cells [12]. This is consistent with literature reports on GR targeting being effective in eliminating LSCs [10,66]. Our data suggest that the PS127-family compounds, and likely 125_1, induce selective AML cytotoxicity by increasing ROS, likely by inhibiting GR, resulting in activation of lethal mitophagy (**Figure 6G**). Additionally, we attempted to generate DOX-resistant MOLM-13 clones through low-dose exposure to evaluate the potential of our lead compounds in a resistant state. However, establishing a stable, highly-resistant population was challenging; cells remained sensitive to redox disruption, as indicated by low cellular viability. Characterizing the ability of PS127-family compounds to maintain efficacy across diverse leukemic states represents an important direction for future investigation.

We did not observe overt adverse effects on viability or GR activity when PBMCs were treated with PS127-family compounds or 125_1 (**Table 1**, **Figure 3G**), suggesting that they do not significantly impact essential housekeeping pathways. Although our study indicates that GR disruption is likely to be the primary driver of cytotoxicity, lead compounds potentially have broader cellular interactions. To investigate this further, RNA-sequencing and proteomic profiling should be performed.

We also evaluated whether our compounds synergize with current AML treatments, as cancer therapy often involves combinations of drugs, like high-dose ara-C and an anthracycline antibiotic (e.g. doxorubicin, daunorubicin, or idarubicin) [3]. Since neither of these compound groups target GR, we tested both ara-C and DOX for synergy with PS127-family compounds. We also tested the tyrosine kinase receptor inhibitor midostaurin, which is often administered to AML patients with FLT3 mutations—about 30% of the patient population [67]. Our compounds showed strong synergy with midostaurin in cells with (MOLM-13 and MV4;11) and without (OCI-AML2) FLT3 mutations, which is a promising result.

While our compounds were also synergistic with DOX, the magnitude was notably lower. This may result from a cascade of effects arising from DOX’s role in inhibiting DNA replication, including interfering with mitochondrial bioenergetics and respiration; this may create mechanistic overlap between DOX and PS127-compounds [68]. Fortunately, PS127-family compounds, unlike DOX, exhibited minimal cardiotoxicity at concentrations that killed AML cells. This suggests that they may have significant promise in replacing DOX in some drug combinations for appropriate patients, including those with acute lymphoblastic leukemia [69]. Our compounds were also synergistic with first and second-generation FLT3 inhibitors, indicating their potential for use in a multitude of scenarios (refractory, relapse) [70]. To further support this notion, our compounds were active in AML patient cells regardless of FLT3 mutation status, a promising result for pre-clinical animal models and one that reinforces that these compounds can target the heterogeneous nature of AML. Additionally, AML cells adapt their proteolytic systems to attenuate cytotoxicity due to misfolded protein stress. This adaptation indicates that autophagy depends on the cell’s proteostatic capacity [71]. To test this, combining proteasome inhibitors and late-stage autophagy inhibitor represents a promising strategy for future studies.

It is widely recognized that personalized therapy is likely to be the next significant advance in oncology; a wider variety of drugs with new mechanisms will facilitate this era of treatment, but more thorough study in a broad spectrum of cancer genotypes will be a necessary pre-requisite for this approach. Overall, in addition to demonstrating a novel therapeutic approach, the molecules identified in this study have provided leads for the development of compounds with the necessary properties to become therapeutics.

## ACKNOWLEDGEMENTS

The research in Kirienko lab was supported by NIH NIGMS (R35GM129294), NCI (R21CA280500), and CPRIT HIHR RP250573 grants to NVK, an NIH NCI supplement (3R21CA280500-01A1S1) to LA, and an NIH NIGMS (T32GM139801) grant to MRD. We thank GCC TIPS for providing support for this project. We thank Dr. Katelyn Baumer (Rice University) for guidance on DSF experiments. *In silico* analysis was performed within the framework of the Program for Basic Research in the Russian Federation (2021-2030) (project 122030100170-5, VVP and LAS). AML patient samples were kindly provided by MD Anderson Cancer Center Leukemia Sample Bank (Director: Prof. Steven Kornblau). Cardiomyocyte cell line was provided by Dr. Natalia Baran.

## AUTHOR CONTRIBUTIONS

MRD and BM conducted the majority of the experiments, performed data analysis and visualization, generated all the figures, drafted the manuscript and contributed to its revision. LA performed experiments, data analysis, and visualization crucial for the review process, and helped draft and edit the manuscript. ET and AVR contributed to the initial *in silico* screen and preliminary experimental work. LAS and VVP performed *in silico* analysis. SRG contributed to compound selection from the *in silico* screen and provided medicinal chemistry expertise on molecules’ potential. NB provided cell lines and methodological expertise and contributed to revising and editing manuscript. NT contributed to informative experiments not included in the manuscript and provide guidance on experiments with stem cells. NVK provided overall funding, crucial support and guidance for this study. NVK also contributed to data analysis, experimental design, and manuscript writing and editing. All authors reviewed the manuscript before submission.

## COMPETING INTERESTS

The authors declare no competing interests.

## MATERIALS AND CORRESPONDENCE

Correspondence and requests for materials should be addressed to the corresponding author.

## DATA AVAILABILITY

The authors declare that the data supporting the findings of this study are available within the paper and its supplementary information files.

## CODE AVAILABILITY

All custom code used to generate figures and statistical analyses is available from the corresponding author upon reasonable request.

---

**Supplementary Figure 1. MDS clustering of screened compounds for apoptosis induction and T/GR inhibition.** Clustering of 161 primary screen hits using MDS algorithm. Two large clusters, 20 and 103, were identified. Cluster 20 contained PS127-family molecules.

**Supplementary Figure 2. Structural similarity of compounds selected for wet-lab validation.** Clustering is based on Tanimoto coefficient generated using PubChem algorithm. Compounds with similarity coefficients at or above 0.75 were connected by line and placed into the same cluster.

**Supplementary Figure 3. Dose-dependent cytotoxic concentration of lead compounds in FLT3-ITD AML cells. A-C**) Dose-response curve of lead compounds in MOLM-13 (A), MV4;11 (B), and PBMCs (C). Cells were treated with a range of concentrations of compound for 72 hours. **D**) Summary of CC_50_ values (µM) derived from dose-response curve. Ratio of CC_50_ in PBMC to the respective leukemic cells was calculated to show selectivity. Error bars show SEMs from three biological replicates.

**Supplementary Figure 4. Synergistic cytotoxicity of PS127_4 in combination with MID and DOX. A**) Viability of MOLM-13 cells (pink bars) and healthy PBMCs (grey bars) following treatment with PS127_4, MID, or their combination. **B**) Viability of MOLM-13 cells and healthy PBMCs following treatment with PS127_4, DOX, or their combination. Error bars show SEM from three biological replicates. Black asterisks indicate comparison of MOLM-13 cells vs. healthy PBMCs under the same combinatorial treatment conditions. Pink asterisks indicate significantly lower survival under combinatorial treatment compared to single PS127-family compound treatment; Blue asterisks indicate significantly lower survival under combinatorial treatment compared to a single MID treatment. Significance was assessed via Student’s *t*-test.

**Supplementary Figure 5. Synergistic effects of PS127-family compounds with commercial chemotherapeutics in MOLM-13.** Representative synergy landscape of MOLM-13 cells (left) or PBMCs (right) with PS127E, PS127_4 or PS127_10 with MID (A, B), DOX (C, D), and ara-C (E-G). Black dots represent the maximum synergy point in MOLM-13 and its correspondence in PBMCs.

**Supplementary Figure 6. Synergistic effect of PS127-family compounds with MID in other AML cell lines.** Representative synergy landscape of MV4;11 with PS127E or PS127_10 and MID (A,B). Representative synergy landscape of OCI-AML2 with lead PS127-family compounds (C-E). Black dots represent the highest synergy score in AML cell lines (right) and its corresponding in PBMCs (left).

**Supplementary Figure 7. Synergistic effect of PS127-family compounds with other FLT3 inhibitors in leukemic cell lines.** Representative synergy landscape of PS127-family compounds and QUIZ in MOLM-13 (A, C) or MV4;11 (B, D). Representative synergy landscape of PS127-family compounds and SORA in MOLM-13 (E, G) or MV4;11 (F, H). Representative synergy landscape of PS127-family compounds and GIL in MOLM-13 (I, K) or MV4;11 (J, L). Black dot represents the highest BLISS score in leukemic cell lines (left) and its correspondence in PBMCs (right).

**Supplementary Figure 8. Synergistic effect of PS127-family compounds with VEN in MOLM-13 cell lines.** Representative synergy landscape of PS127E (A) or PS127_10 (B) and VEN. Black dot represents the highest BLISS score in MOLM-13 cell lines (left) and its correspondence in PBMCs (right).

**Supplementary Figure 9. Antagonistic effect of PS127-family compounds with various autophagy inhibitors in MOLM-13 cell lines.** Representative synergy landscape of lead PS127-family compounds with Wortmannin (A), CQ (B), 3-MA (C), and HCQ (D). Minimum Bliss scores are shown for each combination.

**Supplementary Figure 10. Cytotoxicity of various redox metabolites after 72-hour treatment. A**) Neither GSH (yellow, n=3) nor GSSG (green, n=3) were toxic to AML cells. **B**) Tocopherol (purple, n=3) and **C**) ascorbate (green, n=3) showed cytotoxicity at high concentrations. Cell viability was normalized to solvent-treated as negative control. The highest concentration of tocopherol or ascorbate with minimal toxicity was used to assess their impact on compound-induced cytotoxicity. Bar graphs represent the mean normalized viability from three biological replicates, error bars indicate SEM.

**Supplementary Figure 11. Evaluation of PASS-predicted activities in other top PS127-family compounds and their mechanistic activity to induce cell death. A**) The effect of redox-active compounds (GSH (n = 3), GSSG (n = 3), and α -tocopherol (n = 3) on PS127-induced cytotoxicity in MOLM-13. **B**) The effect of ascorbate or tocopherol supplementation either individually or in combination on cell viability treated with lead PS127-family compounds, **C-D**) Representative histograms of DHE^+^ population upon PS127-family treatment in the presence or in absence of redox metabolites. **E-F**) Representative histogram of MitoSOX Red^+^ population upon PS127-family treatment in the presence or in absence of redox metabolites. Peaks with dashed lines display control treatment or in absence of PS127-molecule, while solid lines represent treatment with PS127-molecule. **G)** Quantification of total ROS upon PS127-family treatment in the presence of glutathione redox metabolites. ROS level was indicated by MFI of DHE (n = 3). **H**) Quantification of mitochondrial ROS level in the presence of redox metabolites. ROS level was indicated by MFI of MitoSOX Red (n = 3). Biological replicates are shown by data points. Statistical significance of apoptotic cells and ROS level was assessed via ANOVA using *post hoc* Dunnett’s test, comparing each treatment against the no redox metabolite group.

**Supplementary Figure 12. Basal OCR of PBMCs and various leukemic cells.** OCR was normalised to the number of live cells (in millions) per chamber volume (n = 4). OCR of leukemic cells were statistically compared to PBMCs using Dunnett’s test.

**Supplementary Table 1. Sample information of primary AML patients.**

**Supplementary Table 2. Sample information of CD34^+^ samples.**

**Supplementary Table 3. Structures of tested compounds from first *in silico* screen selection criteria.**

**Supplementary Table 4. Pairwise Tanimoto coefficient matrix of representative compounds with three predicted activities**

**Supplementary Table 5. Structures of *in silico* hits from non-PS127-family clusters**

## Notes

### Competing Interest Statement

The authors have declared no competing interest.

### Summary of Updates

This version of the manuscript contains additional experiments (panels were added to Main and Supplementary figures) as well as textual changes.

