## Supplementary Figures and Tables for "Cheminformatic identification of small molecules targeting acute myeloid leukemia"

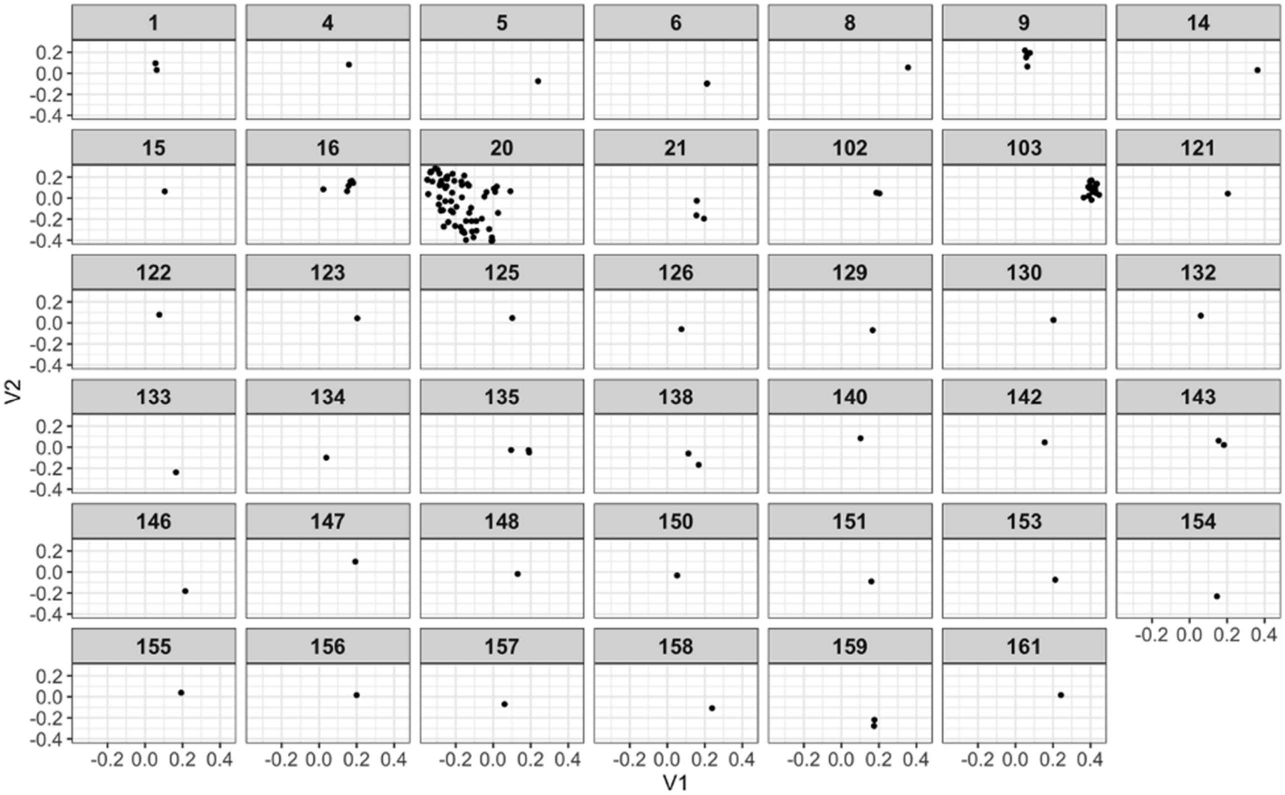


**Supplementary Figure 1.**

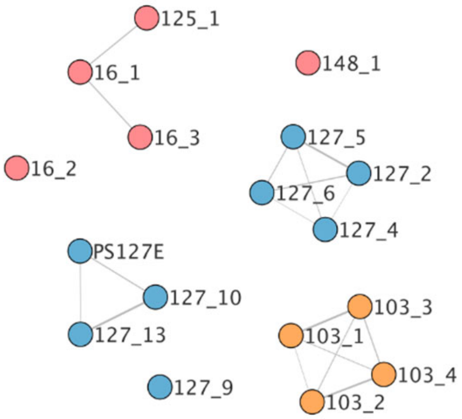


**Supplementary Figure 2.**

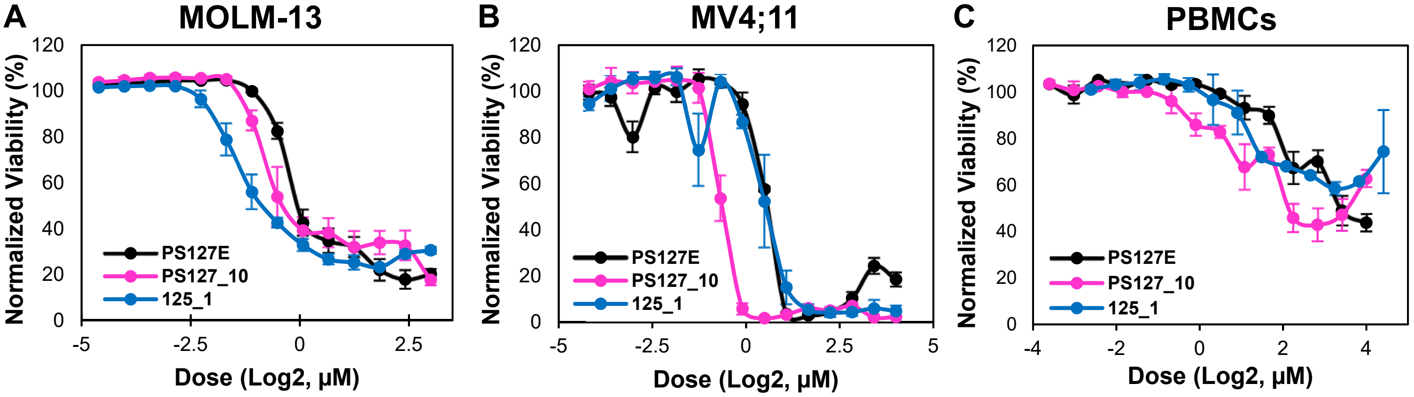


D

CC$_{50}$ ± SEM (µM)

| Compounds | MOLM-13 | MV4;11 | PBMCs | PBMCs/ MOLM-13 | PBMCs/ MV4;11 |
|---|---|---|---|---|---|
| PS127E | 0.88 ± 0.04 | 1.42 ± 0.01 | 11.34 ± 1.94 | 12.89 | 7.99 |
| PS127_10 | 0.63 ± 0.06 | 0.63 ± 0.03 | 8.59 ± 2.42 | 13.63 | 13.63 |
| 125_1 | 0.41 ± 0.04 | 1.45 ± 0.17 | 15.68 ± 0.75 | 38.24 | 10.81 |

**Supplementary Figure 3.**

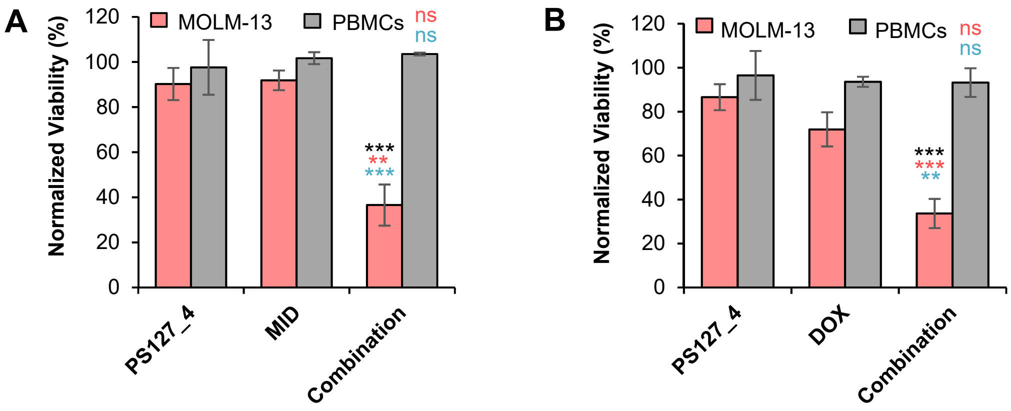


**Supplementary Figure 4.**

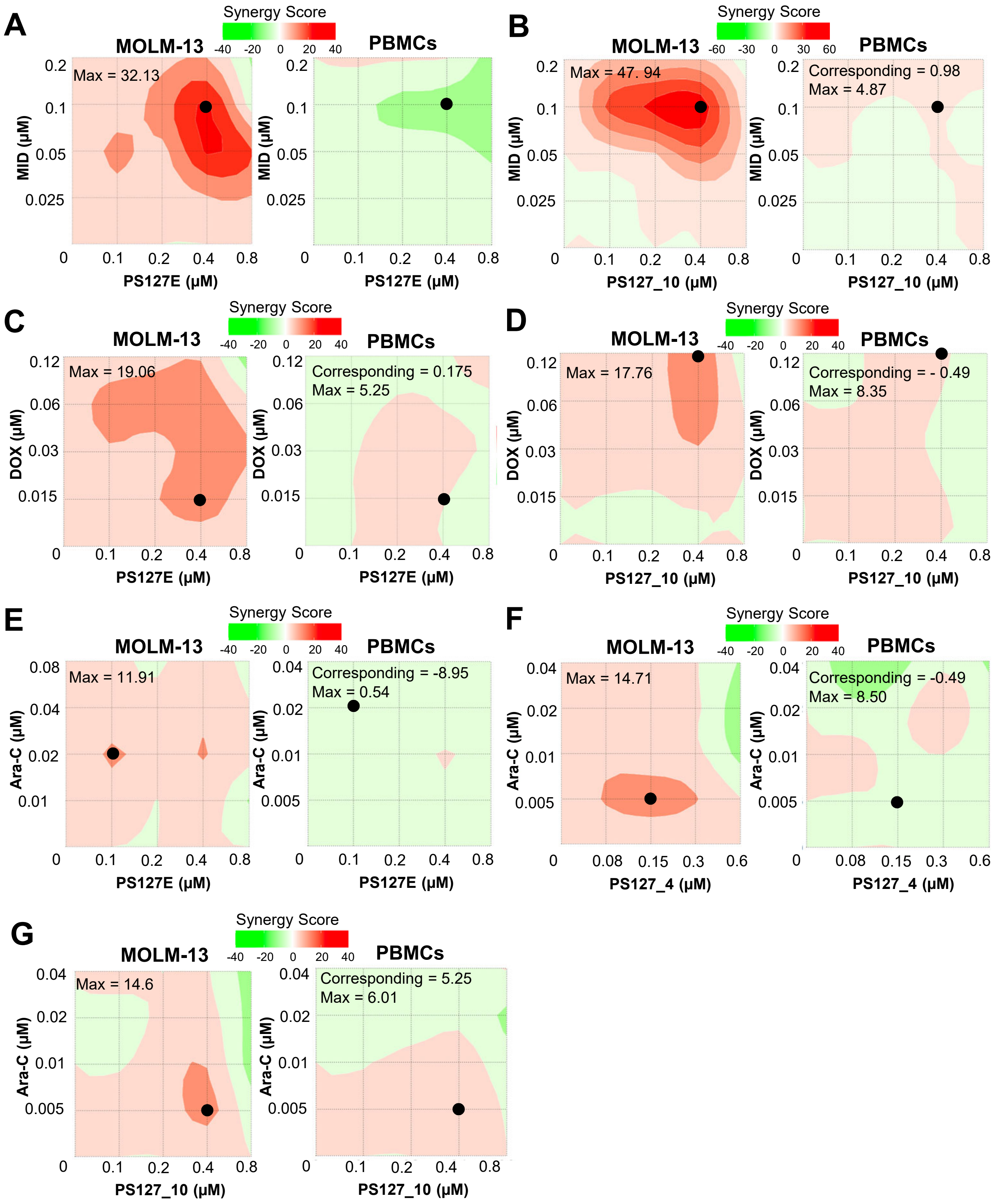


**Supplementary Figure 5.**

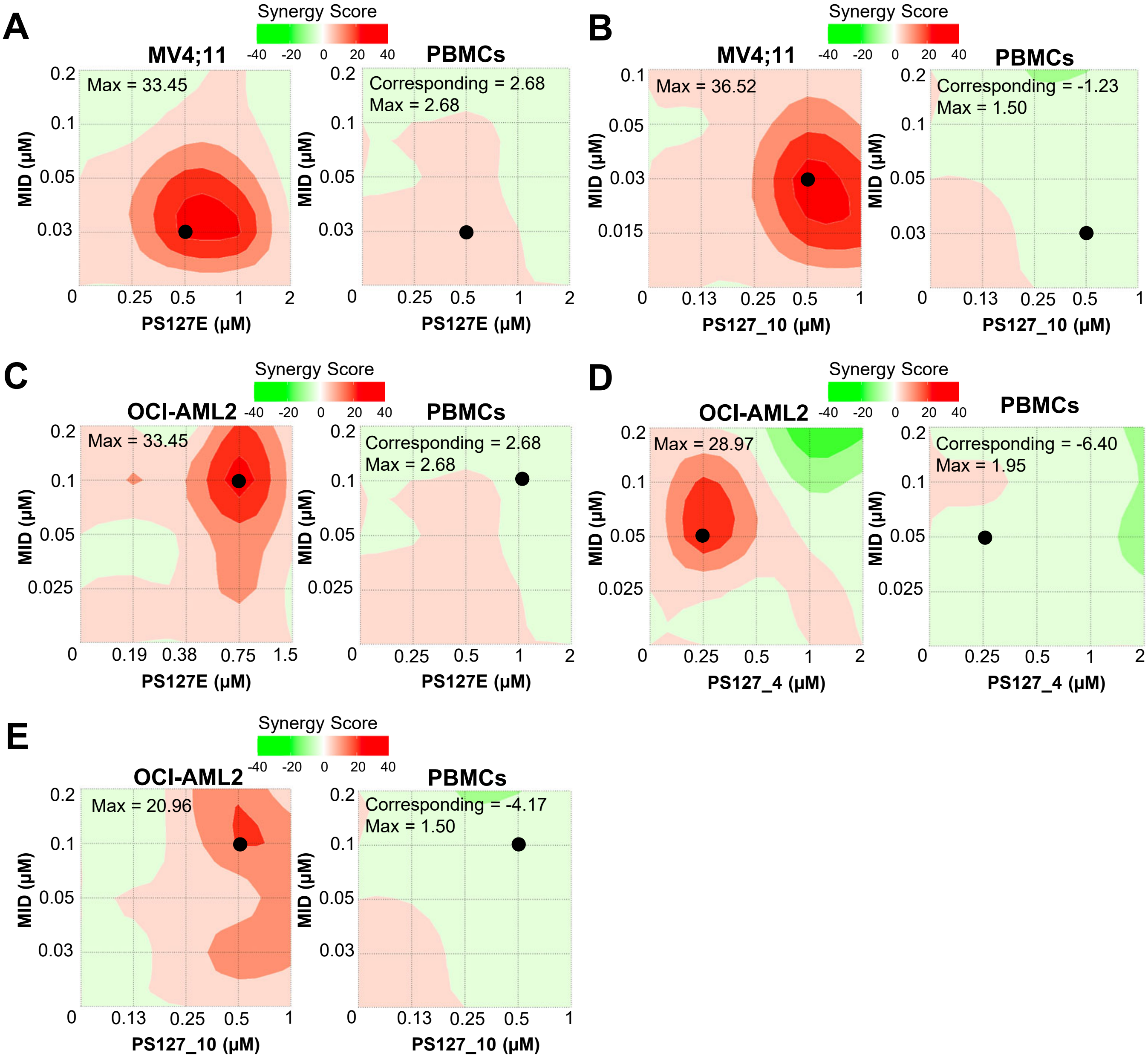


**Supplementary Figure 6.**

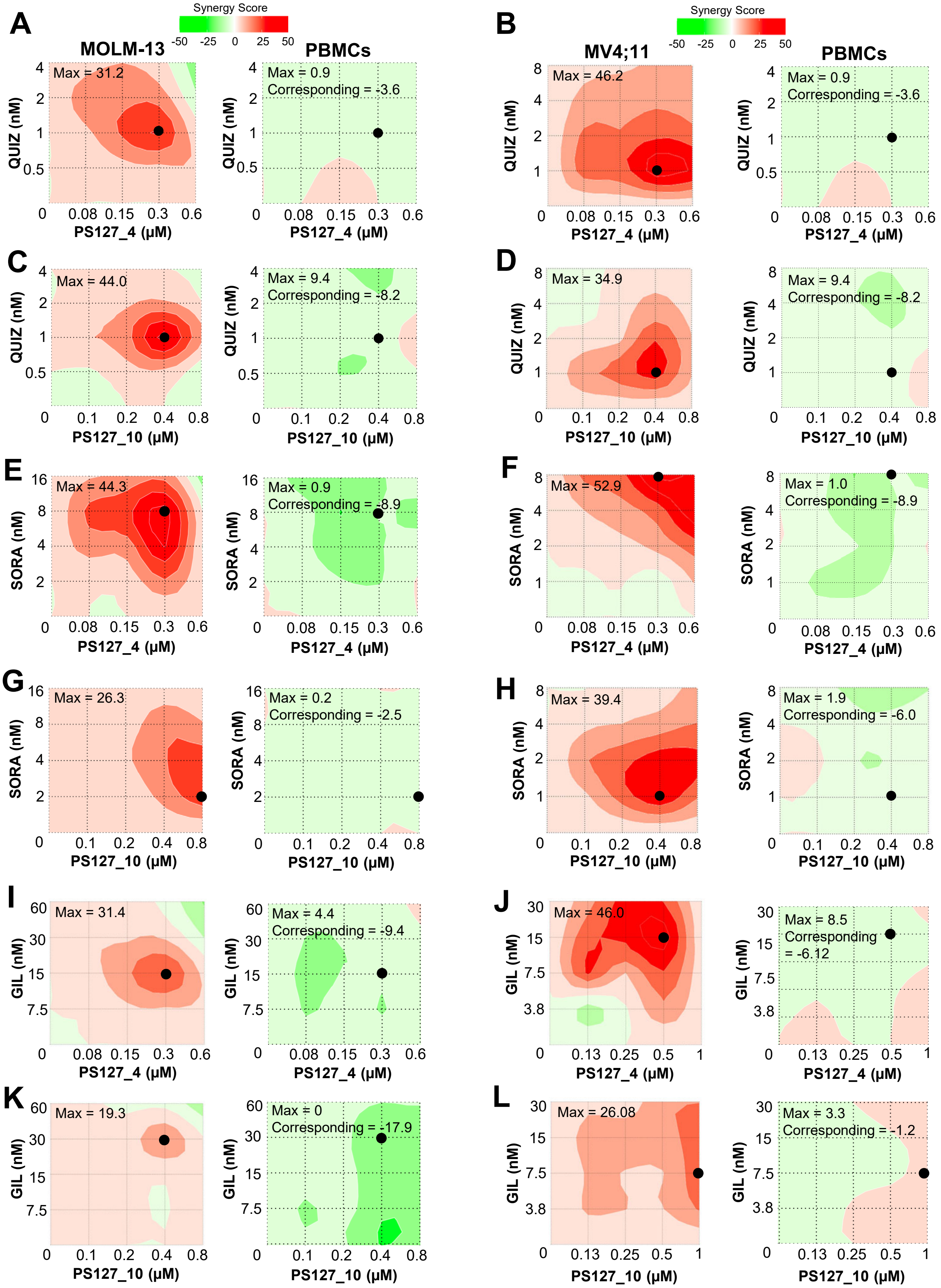


**Supplementary Figure 7.**

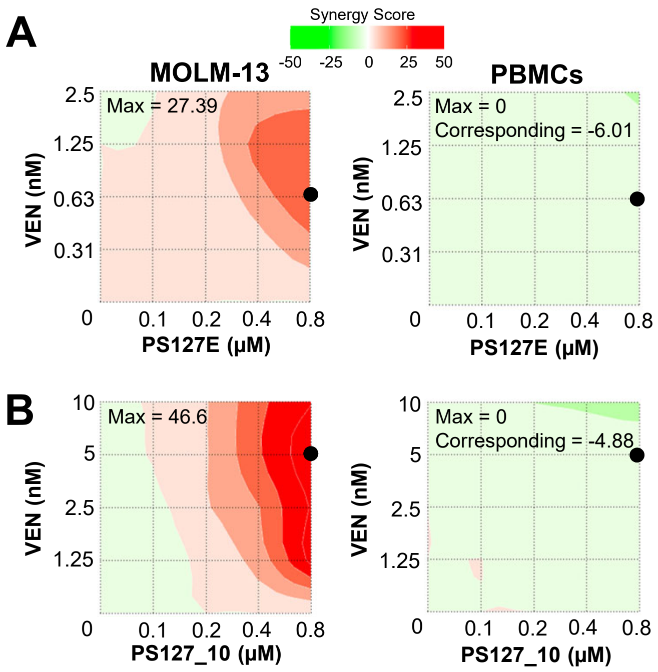


**Supplementary Figure 8.**

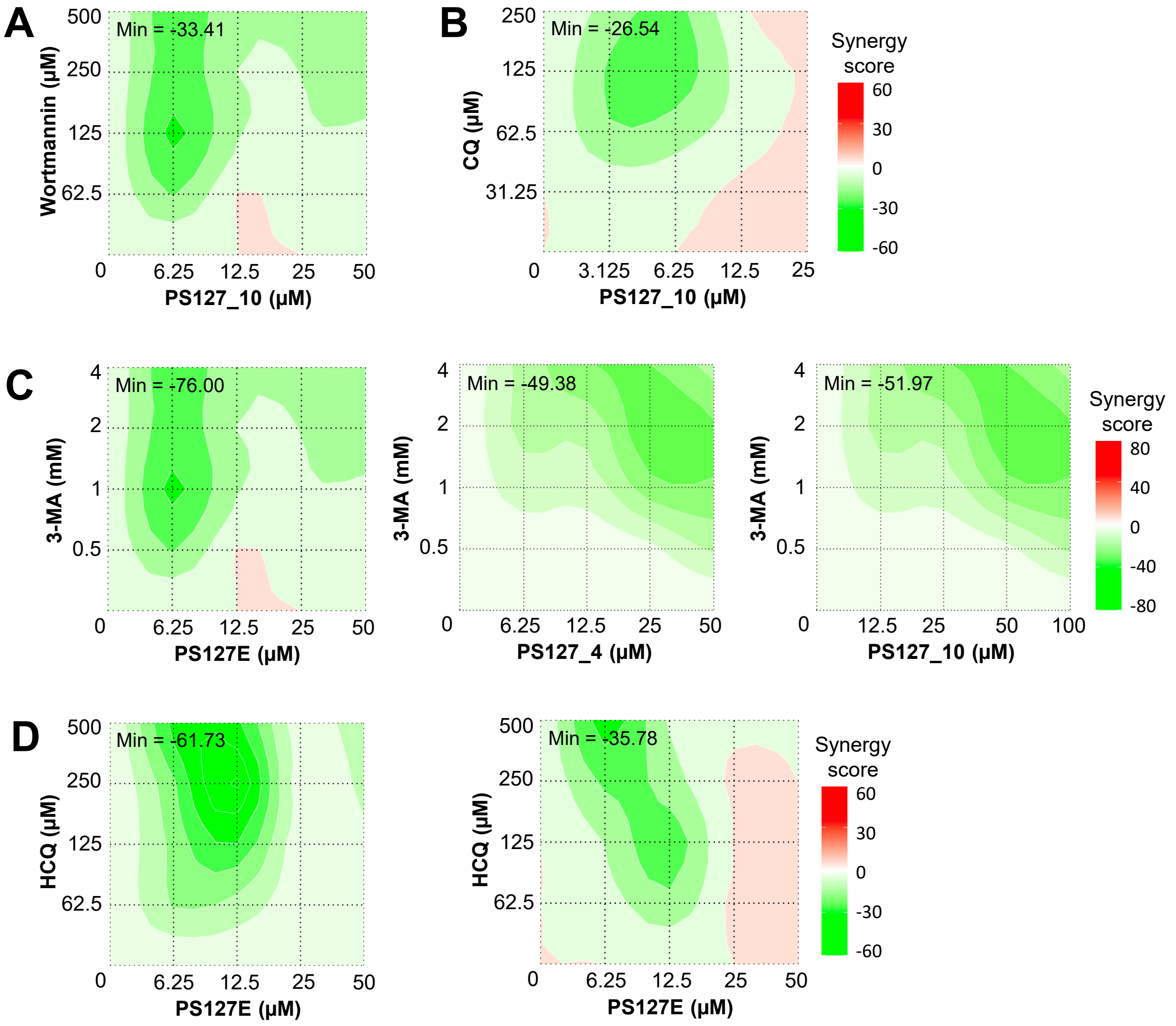


**Supplementary Figure 9.**

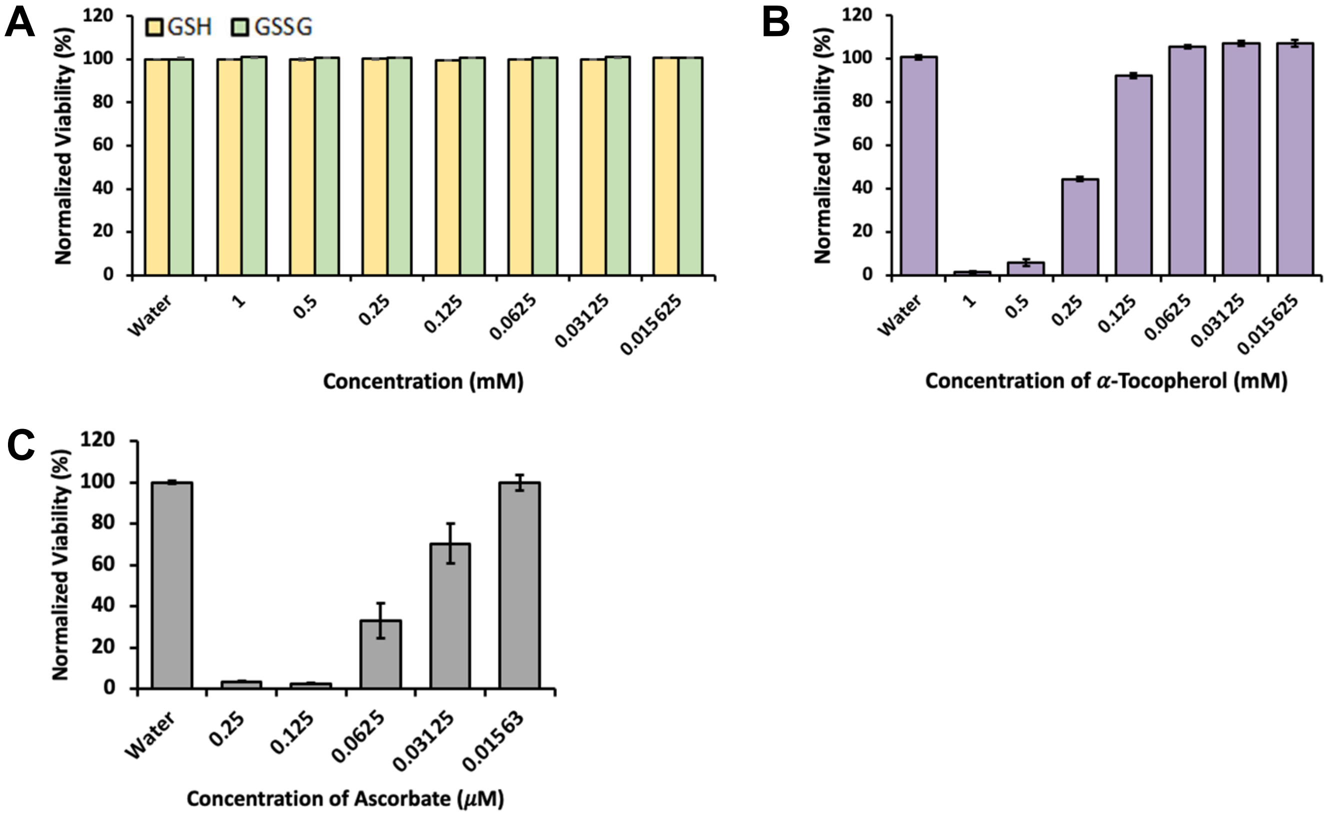


**Supplementary Figure 10.**

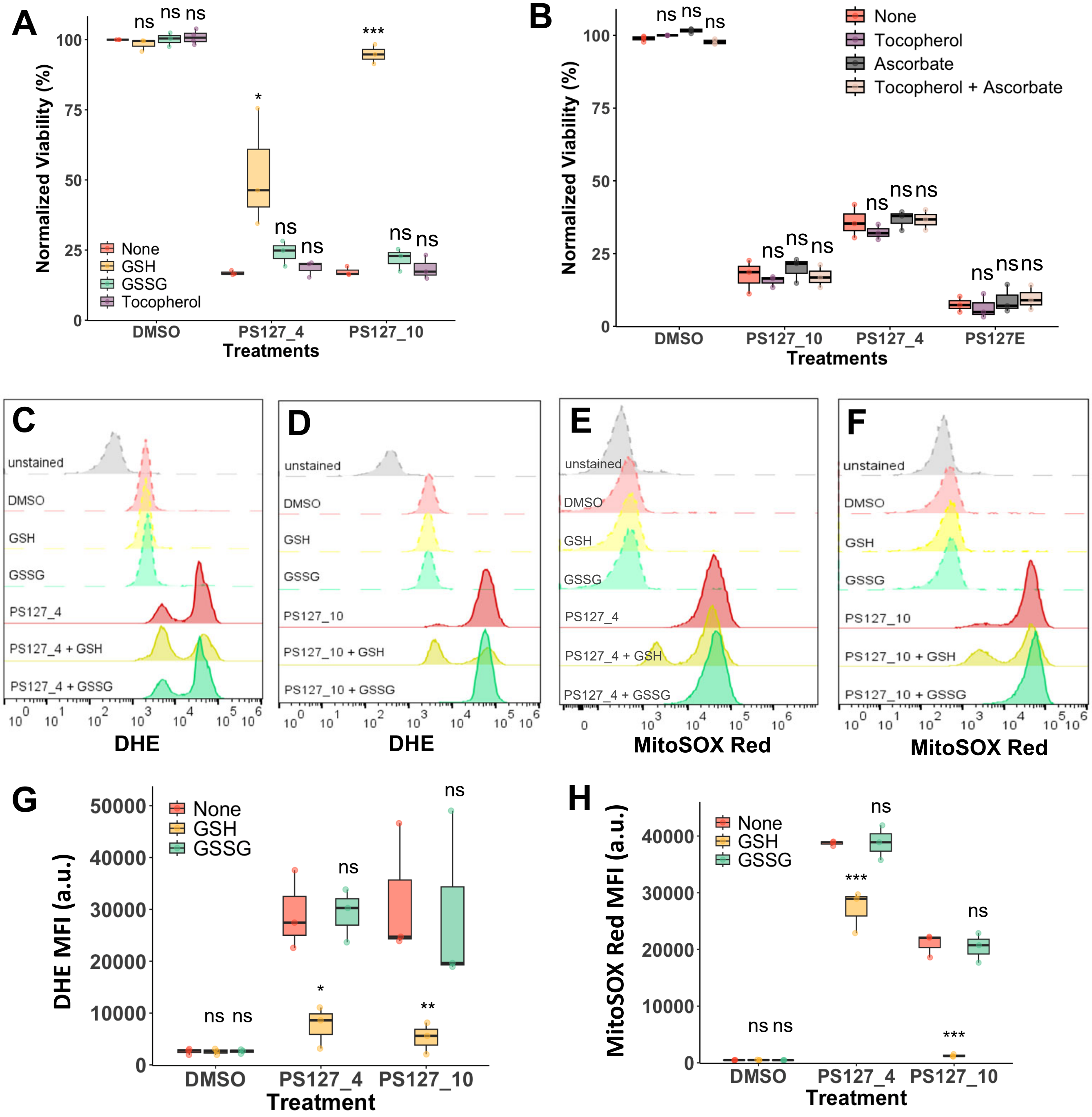


**Supplementary Figure 11.**

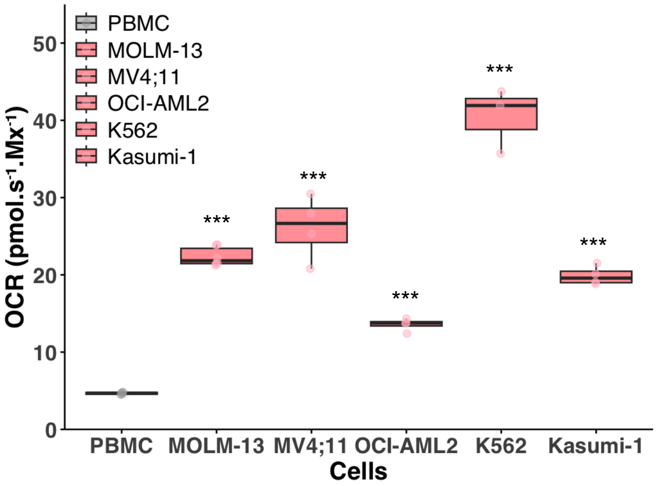


**Supplementary Figure 12.**

**Supplementary Table 1.**

| Patient ID | Cohort | Age | Organ | Cell Count ($x10^6$) | Blast Count ($x10^6$) | WBC ($x10^6$) | Cytogenetics |
|---|---|---|---|---|---|---|---|
| AML-1 | FLT3+ | 60 | Bone Marrow | 13.7 | | | |
| AML-2 | FLT3+ | 46 | Bone Marrow | 14.5 | | | |
| AML-3 | FLT3+ | 58 | Blood | 4 | 31 | 2.8 | 46,XY,-15,+mar[1]/46,XY[19] |
| AML-4 | FLT3+ | 59 | Blood | 4 | 13 | 1.7 | 46,XX[20] |
| AML-5 | FLT3+ | 65 | Blood | 4 | 8 | 1.7 | 42,XY,-6,-11,-12,-16,-17,+mar[1]/46,XY[19] |
| AML-6 | FLT3+ | 67 | Blood | 11 | 26 | 18 | 46,XX,inv(16)(p13.1q22)[20] |
| AML-7 | FLT3+ | 66 | Blood | 10 | 70 | 18.6 | 46,XY[20] |
| AML-8 | FLT3+ | 52 | Blood | 15 | 30 | 10.5 | 46,XY[20] |
| AML-9 | FLT3+ | 68 | Blood | 20 | 89 | 77.3 | 47,XY,+8[3]/49,idem,+11,+13[7]/46,XY,+1,der(1;14)(q10;q10)[3]/46,XY,+1,der(1;13)(q10;q10)[2]/46,XY,+1,der(1;15)(q10;q10)[2]/47,XY,+mar[1]/46,XY[2] |
| AML-10 | FLT3+ | 76 | Blood | 20 | 39 | 45.1 | 47,XY,+mar[1]/46,XY[19] |
| AML-11 | FLT3- | 89 | Bone Marrow | 10 | | | |
| AML-12 | FLT3- | 73 | Bone Marrow | 4.94 | | | |
| AML-14 | FLT3- | 74 | Bone marrow | 5.50 | 54 | 33.9 | 46,XY,del(5)(q11.2)[1]/46,idem,add(12)(p13)[16]/46,idem,der(7)add(7)(p13)add(7)(q32),add(12)(p13)[3] |
| AML-15 | FLT3- | 68 | Bone marrow | 15 | 21 | 3.7 | 46,XX[20] |
| AML-16 | FLT3- | 76 | Blood | 20 | 40 | 15.8 | 45,X,-Y[14]/46\R\49,XY,+Y,+Y,+13,i(17)(q10)[cp6] |
| AML-17 | FLT3- | 78 | Blood | 20 | 95 | 23.7 | 46,XX[20] |

**Supplementary Table 2.**

| Donor ID | Gender | Age | Cell count ($x10^5$) | Organ | CD34 | Purity (%) |
|---|---|---|---|---|---|---|
| Primary-1 | Male | 21 | 8 | Bone marrow | + | 98 |
| Primary-2 | Male | 21 | 7 | Bone marrow | + | 96 |
| Primary-3 | Male | 25 | 9 | Bone marrow | + | 92 |

**Supplementary Table 3.**

| No | PS compound | MolPort ID | MW (g/mol) | Structure | $CC_{50}$, μM, MOLM-13 |
|---|---|---|---|---|---|
| 1. | 127_1 | Molport-002-717-654 | 150.137 | 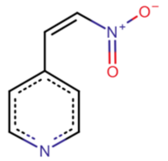 | >8 |
| 2. | 127_2 | Molport-019-923-348 | 185.13 | 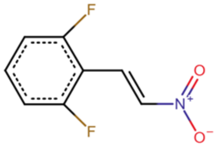 | 1.62 |
| 3. | 127_3 | Molport-009-653-856 | 193.158 | 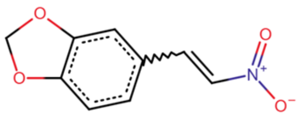 | 2.41 |
| 4. | 127_4 | Molport-002-918-210 | 249.269 | 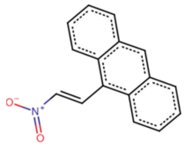 | 0.33 |
| 5. | 127_5 | Molport-002-918-866 | 217.147 | 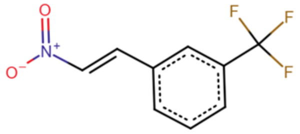 | 1.15 |
| 6. | 127_6 | Molport-023-299-362 | 174.159 | 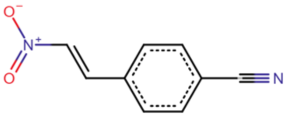 | 1.05 |
| 7. | 127_8 | Molport-002-951-542 | 192.218 | 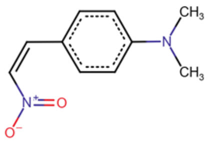 | 6.03 |
| 8. | 127_9 | Molport-030-040-762 | 271.114 | 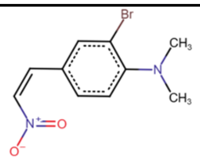 | 0.59 |
| 9. | 127_10 | Molport-003-827-524 | 259.261 | 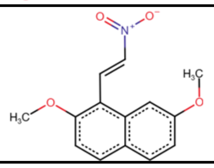 | 0.63 |
| 10. | 127_13 | Molport-003-826-996 | 287.315 | 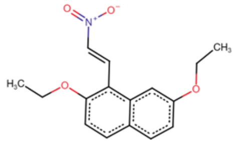 | 2.15 |

| No | PS compound | MolPort ID | MW (g/mol) | Structure | $CC_{50}$, µM, MOLM-13 |
|---|---|---|---|---|---|
| 11. | 127_14 | Molport-019-686-955 | 220.272 | 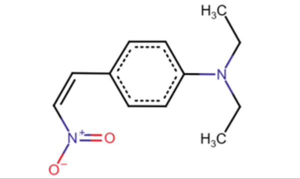 | 3.68 |
| 12. | 127_20 | Molport-000-153-349 | 183.59 | 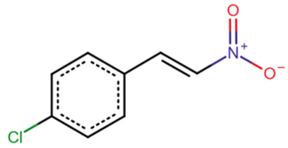 | 1.41 |
| 13. | 127_21 | Molport-002-918-274 | 254.198 | 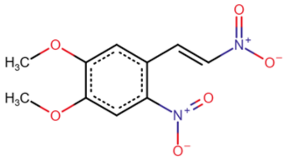 | >8 |
| 14. | 127E | Molport-000-156-857 | 179.175 | 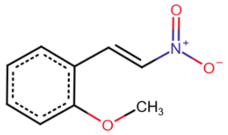 | 0.88 |
| **Other Tested Compounds** | | | | | |
| 15. | 103_1 | Molport-003-185-960 | 343.335 | 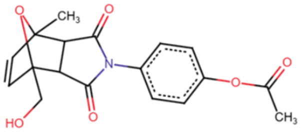 | >10 |
| 16. | 103_2 | MolPort-001-002-648 | 321.332 | 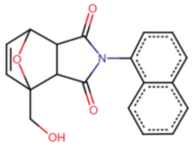 | >10 |
| 17. | 103_3 | MolPort-003-874-156 | 301.298 | 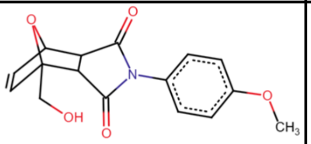 | >10 |
| 18. | 103_4 | MolPort-002-807-280 | 299.326 | 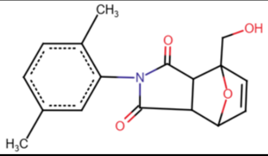 | >10 |

### Supplementary Table 4.

| | 103_1 | 103_2 | 103_3 | 103_4 | PS127_2 | PS127_4 | PS127_5 | PS127_6 | PS127_9 | PS127_10 | PS127_13 | PS127E | 16_1 | 16_2 | 16_3 | 125_1 | 148_1 |
|---|---|---|---|---|---|---|---|---|---|---|---|---|---|---|---|---|---|
| **103_1** | | | | | | | | | | | | | | | | | |
| **103_2** | 0.764 | | | | | | | | | | | | | | | | |
| **103_3** | 0.944 | 0.807 | | | | | | | | | | | | | | | |
| **103_4** | 0.799 | 0.928 | 0.833 | | | | | | | | | | | | | | |
| **PS127_2** | 0.314 | 0.360 | 0.327 | 0.367 | | | | | | | | | | | | | |
| **PS127_4** | 0.312 | 0.412 | 0.324 | 0.377 | 0.753 | | | | | | | | | | | | |
| **PS127_5** | 0.309 | 0.354 | 0.322 | 0.360 | 0.933 | 0.787 | | | | | | | | | | | |
| **PS127_6** | 0.307 | 0.352 | 0.320 | 0.375 | 0.848 | 0.758 | 0.817 | | | | | | | | | | |
| **PS127_9** | 0.415 | 0.457 | 0.431 | 0.474 | 0.604 | 0.548 | 0.588 | 0.596 | | | | | | | | | |
| **PS127_10** | 0.476 | 0.449 | 0.486 | 0.410 | 0.540 | 0.627 | 0.565 | 0.523 | 0.437 | | | | | | | | |
| **PS127_13** | 0.508 | 0.457 | 0.495 | 0.426 | 0.515 | 0.611 | 0.538 | 0.500 | 0.421 | 0.952 | | | | | | | |
| **PS127E** | 0.489 | 0.409 | 0.509 | 0.424 | 0.644 | 0.581 | 0.626 | 0.620 | 0.500 | 0.833 | 0.794 | | | | | | |
| **16_1** | 0.278 | 0.346 | 0.290 | 0.327 | 0.593 | 0.573 | 0.573 | 0.567 | 0.430 | 0.420 | 0.401 | 0.456 | | | | | |
| **16_2** | 0.376 | 0.450 | 0.391 | 0.432 | 0.615 | 0.596 | 0.598 | 0.593 | 0.659 | 0.456 | 0.439 | 0.492 | 0.680 | | | | |
| **16_3** | 0.279 | 0.368 | 0.290 | 0.341 | 0.537 | 0.684 | 0.568 | 0.546 | 0.411 | 0.489 | 0.467 | 0.434 | 0.831 | 0.593 | | | |
| **125_1** | 0.305 | 0.342 | 0.317 | 0.357 | 0.635 | 0.561 | 0.614 | 0.663 | 0.471 | 0.424 | 0.406 | 0.500 | 0.836 | 0.634 | 0.744 | | |
| **148_1** | 0.434 | 0.397 | 0.443 | 0.403 | 0.466 | 0.417 | 0.455 | 0.513 | 0.366 | 0.574 | 0.551 | 0.578 | 0.424 | 0.382 | 0.394 | 0.491 | |

**Supplementary Table 5.**

| No | Compound | MolPort ID | MW (g/mol) | Structure | $CC_{50}$, μM, MOLM-13 |
|---|---|---|---|---|---|
| 19. | 16_1 | Molport-020-171-975 | 244.37 | 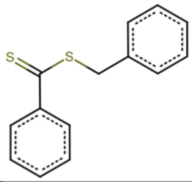 | ND, <10 |
| 20. | 16-2 | Molport-003-005-421 | 289.37 | 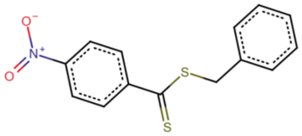 | ND, <10 |
| 21. | 16-3 | MolPort-003-800-321 | 218.33 | 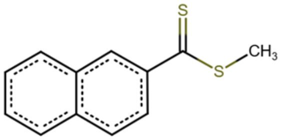 | ND, >10 |
| 22. | 125_1 | MolPort-006-727-986 | 221.34 | 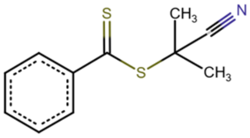 | 0.41 |
| 23. | 148_1 | MolPort-003-850-809 | 202.169 | 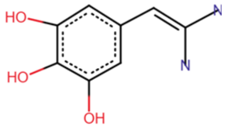 | ND, >10 |

ND = not determined
