## Supplementary Methods for "Cheminformatic identification of small molecules targeting acute myeloid leukemia"

### SUPPLEMENTAL METHODS

#### SM1 Cell culturing of AML cell lines, PBMCs, cardiomyocytes, AML patient samples, and $CD34^+$ progenitor cells

AML cell cultures were maintained in standard culture media (RPMI-1640 medium supplemented with 2 mM L-glutamine and sodium bicarbonate (Sigma-Aldrich), 10% Corning™ Regular Fetal Bovine Serum (FBS) (Heat Inactivated) and 1% penicillin-streptomycin solution (P/S) (Sigma-Aldrich)) in a 37°C humidified atmosphere with 5% $CO_2$. MOLM-13 was used as an AML cell line model, unless specified otherwise. H9C2 cardiomyocytes were subcultured in similar condition via trypsinization at 70% confluence prior to experimental treatment. A new tube was thawed after about 50 passages.

Peripheral blood mononuclear cells (PBMCs) were obtained from the Gulf Coast Regional Blood Centre (Houston, TX, USA). PBMCs were isolated using Ficoll®-Paque PREMIUM (Cytiva) according to manufacturer's instructions. PBMCs were maintained overnight (at least 16 hours) at a minimum density of $10^6$ cells/mL before experimental use.

Cryopreserved AML patient samples were anonymized to maintain patient confidentiality **(Supplementary Table 1)**. Frozen cells were thawed and transferred into thawing media (RPMI-1640 medium with 20% FBS, 1 mM $MgSO_4$, and 1 mM EDTA). The vials were washed twice with thawing media and spun down for 5 minutes at 400 g. The samples were washed with RPMI-1640 medium (10% FBS, 1% P/S) and spun down twice more. Samples were then cultured in 20% FBS, 1% P/S RPMI-1640 at a $10^6$ cells/mL the day prior to experimental set-up.

All assays with patient samples used patient-experimental media (RPMI-1640, 2.5% FBS, 1% P/S). AML patient cells were seeded at 30,000 cells per well for synergy experiments with PS127_10 and MID or DOX (depending on sample size), as well as supplementation assays of cell treated with 4 μM of compound and 500 μM of GSH. Experimental plates were imaged after 72 hours. Routine respiration of AML patient cells (pmol·$s^{-1}$·$Mx^{-1}$) was measured in the 0.5 mL chamber of the Oroboros in patient-experimental medium at $20^6$ cells/mL. Routine respiration was also measured for MOLM-13 ($10^6$ cell/mL) and PBMCs ($20^6$ cells/mL) in the 0.5 mL chamber to parallel conditions. Measurements were normalized to the number of live million cells (Mx) per mL.

**SM2 *In silico* screen**

To identify selective cytotoxic against AML cells, we screened a commercially available library of approximately 4.2 million compounds (MolPort [https://www.molport.com/shop/index]) using PASS software version 2022. PASS is a ligand-based cheminformatics approach that is target-agnostic and predicts biological activity solely from chemical structure. Library was filtered for molecules exhibiting three biological activities previously linked to selective cytotoxicity in AML cells: (1) apoptotic agonism, (2) thioredoxin/glutathione reductase (T/GR) inhibition, and (3) autophagic induction. PASS uses a training set comprising thousands of known bioactive compounds to predict over 6,000 types of biological activities based on molecular structure descriptors. For each activity, it calculates two probabilities: Pa (probability of being active, ranging from 0 to 1) and Pi (probability of being inactive, ranging from 0 to 1). In the primary screening, a compound was considered a hit if the difference ΔP (Pa – Pi) for each of the three target functions (apoptotic agonism, T/GR inhibition, and autophagy induction) was ≥ 0.7. This stringent threshold was chosen to minimize false positives.

Because the primary screen alone is typically insufficient, we conducted an additional refinement of the hit list. Based on our previous findings highlighting the critical role of autophagy in the mechanism of PS127-family compounds, we applied a secondary filter requiring a Pa value ≥ 0.7 specifically for the "autophagy induction" activity. This refinement step further reduced the final hit list. The 93 final hits were clustered by structural similarity using the Multidimensional Scaling (MDS) algorithm implemented in ChemMine Tools (https://chemminetools.ucr.edu/), with a similarity cut-off of 0.4 for cluster formation.

To visualize and analyze structural relationships among the hits, pairwise Tanimoto coefficients were calculated using PubChem fingerprints, and the resulting network was visualized in Cytoscape 3.10.3 (https://cytoscape.org/).

Tanimoto coefficients calculated as:

$$T(a,b) = \frac{N_c}{N_a + N_b - N_c}$$

$N_c$ – number of features or descriptors common to both molecules

$N_a$ – number of features or descriptors of the first molecule in the pair

$N_b$ – number of features or descriptors of the second molecule in the pair

To validate the output of the *in silico* screen, a representative subset of compounds was selected for experimental testing. The selection strategy was designed to achieve two primary goals: (1) to sample the chemical space of the most populated cluster, and (2) to test the core hypothesis that the predicted bioactivity profile serves as a generalizable predictor of biological effect, independent of any specific chemical scaffold.

The identified hits were first partitioned into distinct structural classes using molecular similarity-based clustering. From the predominant structural class, which comprised analogs of the initial lead compound PS127E, a representative set of molecules was selected. This selection prioritized maximal structural diversity within the class rather than adherence to a single quantitative metric, thereby ensuring a comprehensive exploration of its chemical space.

To decouple the observed biological effects from the primary chemical scaffold, a separate panel of compounds was chosen from the remaining structural classes. These compounds were intentionally selected for their structural dissimilarity to the main cluster (i.e., their distinct core scaffolds) but were predicted by the PASS algorithm to share the same trio of high-probability biological activities. This design enabled direct testing of whether the predicted functional profile - rather than structural similarity itself - served as the critical determinant of the desired phenotypic outcome.

**SM3 Cell viability assays**

**1. Cytotoxicity assay**

Each *in silico*-identified compounds and commercial chemotherapeutics were prepared as a 10 mM stock solution in DMSO, aliquoted, and stored long-term at -20°C and 4°C for short-term use.  Cells were treated with compounds or solvent controls in experimental media (RPMI-1640 with 1% FBS, 1% P/S) and stained with Hoechst 33342 and propidium iodide (PI) to detect total and dead cells, respectively. Plates were spun down for 5 minutes at 1,000 rcf in the Hermile Z446 Benchmark. Stained cells were then analyzed on a BioTek plate reader (Cytation 5 Cell Imaging Multi-Mode Reader) with Gen5 software.

Cells were seeded in 96-well plates (5,000 AML cells/well or 50,000 PBMCs/well) and immediately treated with screened compound or the solvent control, DMSO in experimental media (RPMI-1640, 1% FBS, 1% P/S). AML patient samples were seeded at 30,000 cells/well in patient-experimental media (RPMI-1640, 2.5% FBS, 1% P/S). Cells were treated with compounds or solvent controls for 72 hours in experimental media. Preliminary cytotoxicity assays assessed compound efficacy at 10 µM of compound.  The concentration of 50% cytotoxicity ($CC_{50}$) for each compound was determined by fitting dose-response models using the dose-response curve (drc) package in R [27]. $CC_{50}$ assays were performed using 2-fold serial dilutions. The highest concentration of solvent control was used, never exceeding 0.5% (v/v). Cell viability was normalized to the solvent-control viability. Three technical replicates were performed per dose with at least three biological replicates.

### 2. **Synergy assay**

Cell viability upon combinatorial treatment was assessed by following previously described guidelines [22]. Commercially available AML chemotherapeutics (Doxorubicin [DOX] [Ark Pharm Inc.], cytarabine [ara-C] [Accela Chembio Inc], midostaurin [MID] [MedChemExpress], or venetoclax [VEN] [ChemieTek]) or other FLT3 inhibitors (gilteritinib [GIL] [Thermo Scientific Chemicals], quizartinib [QUIZ] [Targetmol Chemicals], sorafenib [SORA] [Medchemexpress]) were used.

Cells were seeded in 96-well plates and immediately treated for 72 hours. Single drug treatments and combinations were assessed in technical duplicates for at least three biological replicates. The highest DMSO concentration (not more than 0.5%) was used as the solvent-control. Viability was measured as in cytotoxicity assay to determine synergy for each combinatorial treatment using SynergyFinder package in R (https://synergyfinder.fimm.fi) [28]. Synergy scores were calculated using Bliss model. Bliss synergy scores indicate that scores larger than 10 are synergistic; scores between -10 and 10 are additive, and scores below -10 are antagonistic. We further categorized synergistic strength as follows: low synergy (10-20), medium synergy (>20-30), and high synergy (>30). Synergy plots were created using the same package and visualized in a 2D format. Synergistic potential of combinations was determined by taking the average of maximum synergy (± Standard Error of the Mean (SEM)) scores from at least three biological replicates. Corresponding PBMC synergy matrices were performed to assess potential synergy in healthy cells.

**SM4 Evaluation of T/GR inhibition activity**

**1. Thioredoxin reductase activity assay**

To assess TrxR activity, MOLM-13 cells seeded at density of $10^6$ cells/mL were treated with PS127-compound at $CC_{50,\ 72hours}$ for 48 hours. DMSO was used as solvent-control. Cell lysates were prepared using sonication. TrxR activity was assessed using the Thioredoxin Reductase Colorimetric Assay Kit at 414 nm according to manufacturer's instructions. PS127-treated sample, rat liver TrxR as positive control, and solvent-control were assessed in duplicate.

**2. GSH-to-GSSG ratio assay**

The ratio of GSH to GSSG was assessed using the EnzyChrom™ Glutathione GSH/GSSG Assay Kit. MOLM-13 cells were treated with similar conditions as above. Samples were prepared following manufacturer's instructions for GSSG and total glutathione measurements. The amount of GSSG and total glutathione was determined from $OD_{412nm}$ measurements at 0 and 10 minutes. GSH or GSSG concentration was normalized to total protein concentration, determined by Pierce™ BCA Protein Assay Kit and solvent-control.

**3. Supplementation of exogenous redox metabolites**

GSH (Thermo Scientific Chemicals), GSSG (TCI America™), N-acetylcysteine (NAC)

(Thermo Scientific Chemicals), tocopherol (Thermo Scientific Chemicals), and freshly prepared ascorbate (TCI Chemicals) were dissolved in sterile water. Redox metabolites were added as is into experimental media. MOLM-13 cells were simultaneously treated with redox metabolites and tested compounds at their 4X $CC_{50}$ for 72 hours to assess rescue of cell viability. Cell viability was assessed as per cytotoxicity assay.

**4. GR activity assay**

Cells were treated with lead compounds (2 µM PS127E, 1 µM PS127_10, or 2 µM 125_1) or solvent control for 24 hours. Cells were lysed using sonication in ice and assessed using DetectX® Glutathione Reductase Fluorescence Activity Kit. Human GR was used as a reference. GR activity upon treatment was normalized to total protein concentration measured using Pierce™ BCA Protein Assay Kit.

### SM5 Thermal shift / Differential Scanning Fluorometry assay

DSF was performed by combining 2.5 µM of human GSR (Sigma-Aldrich), 20 µM of compound, and 5X of SYPRO® Orange Protein Gel Stain (ThermoFisher) in 20 mM HEPES, pH 7.4, 100 mM NaCl with 1% DMSO buffer solution [30]. The following controls were used: no protein controls, compound-fluoresce interference, and autofluorescence plate controls. The known GSR inhibitor, 2-AAPA (Sigma-Aldrich), was used as a positive experimental control. Sealed PCR plates were continuously heated with gradual increase in temperature from 25 to 95°C at 1-minute intervals using the PREP BioRad qPCR instrument. RFU was then determined through the FRET channel. Tm was determined by the derivative of relative fluorescence units (dRFU) using the open access tool, DSFworld [31]. The thermal shift analysis was performed with the same materials as described above with a concentration gradient of PS127_10 from 2.5 to 40 µM. Analysis was performed in Thermott, using dRFU as the binding parameters and using their pre-determined thermodynamic parameters for the monomeric oligomer sequence (522 amino acids) [32]. Three technical replicates per four biological replicates were inputted into the binding curve to determine dissociation constant ($K_d$).

### SM6 Bioenergetic measurements

#### 1. ROS and apoptosis evaluation

Total and mitochondrial ROS were measured using dihydroethidium (DHE) (Invitrogen™) or MitoSOX™ Red (Invitrogen™) staining, respectively, as described in manufacturer's protocols. Annexin V-FITC (Thermo Fisher Scientific)/PI/Hoechst staining was used to detect live, apoptotic, and dead cells as previously described [13]. MOLM1-13 cells were seeded at a density of $10^6$ cells/mL and treated with 8 µM of lead compound or 4 µM of PS127_10 for 24 hours. DMSO was used as solvent-control. Flow cytometry was performed on SONY MA900 Cell Sorter. Mean fluorescence intensities (MFIs) were analyzed in FlowJo software 10.8.1.

#### 2. Oxygen Consumption Rate (OCR)

Instrumental background calibration with MiR05 (Oroboros) and oxygen air calibration with experimental RPMI-1640 media were performed according to standardized protocols at 37ºC in the NextGen-O2k instrument (Oroboros Instrument, Innsbruck, Austria).

MOLM-13 cells were treated for 3 hours with 2 µM of compound or DMSO at $10^6$ cells/mL and transferred into the 2-mL Duran® glass chambers maintained at 37°C. Oxygen concentration and negative slope were measured every 2 seconds and stirred at 750 rpm. Cell concentration and viability were measured with the Invitrogen Countess II Automated Cell Counter using Trypan-Blue (0.4%). OCR [pmol·$s^{-1}$·$Mx^{-1}$] was calculated from the average of oxygen negative slope over time (pmol·$s^{-1}$) of a stable 5 min interval (~10 to 15 minutes after experimental start), normalized to the number of live million cells (Mx) per mL.

### 3. ATP measurements

To assess ATP levels, cells were seeded per well into an opaque 96-well plate and incubated with either 2 µM compound or corresponding volume of solvent control (DMSO) for 3 hours at a final volume of 100 µL in experimental RPMI-1640 medium. Each biological replicate was analyzed in three technical replicates. Viability was confirmed to be at least 90% based on Hoechst/PI staining before proceeding with cell lysis. After imaging, 50 µL of media was removed from each well. Subsequently, 50 µL of CellTiter-Glo® 2.0 (Promega) solution was added to each well. Plate was then kept in the dark, gently shaken for 1 minute to induce cell lysis, and incubated at room temperature for 10 minutes to allow luciferase activation. The luciferase signal was measured via luminescence on the BioTek plate reader with Gen5 software. The total ATP production was then calculated by dividing the luminescence signal by the total number of live cells per well.
